# The transcription factor Roc1 is a regulator of cellulose degradation in the wood-decaying mushroom *Schizophyllum commune*

**DOI:** 10.1101/2021.06.08.446897

**Authors:** Ioana M. Marian, Peter Jan Vonk, Ivan D. Valdes, Kerrie Barry, Benedict Bostock, Akiko Carver, Chris Daum, Harry Lerner, Anna Lipzen, Hongjae Park, Margo B. P. Schuller, Martin Tegelaar, Andrew Tritt, Jeremy Schmutz, Jane Grimwood, Luis G. Lugones, In-Geol Choi, Han A. B. Wösten, Igor V. Grigoriev, Robin A. Ohm

**Author notes:** Institute of Hydrobiology, Biology Centre of the Czech Academy of Sciences, České Budějovice, Czech Republic (Hongjae Park).

## Abstract

Wood-decaying fungi of the class Agaricomycetes (phylum Basidiomycota) are saprotrophs that break down lignocellulose and play an important role in the nutrient recycling. They secrete a wide range of extracellular plant cell wall degrading enzymes that break down cellulose, hemicellulose and lignin, the main building blocks of plant biomass. Although the production of these enzymes is regulated mainly at the transcriptional level, no activating regulators have been identified in any wood-decaying fungus in the class Agaricomycetes. We studied the regulation of cellulase expression in the wood-decaying fungus *Schizophyllum commune*. Comparative genomics and transcriptomics on two wild isolates revealed a Zn_2_Cys_6_-type transcription factor gene (*roc1*) that was highly up-regulated during growth on cellulose, when compared to glucose. It is only conserved in the class Agaricomycetes. A *roc1* knockout strain showed an inability to grow on medium with cellulose as sole carbon source, and growth on cellobiose and xylan (other components of wood) was inhibited. Growth on non-wood-related carbon sources was not inhibited. Cellulase activity was reduced in the growth medium of the Δ*roc1* strain. ChIP-Seq identified 1474 binding sites of the Roc1 transcription factor. Promoters of genes involved in lignocellulose degradation were enriched with these binding sites, especially those of LPMO (lytic polysaccharide monooxygenase) CAZymes, indicating that Roc1 directly regulates these genes. A GC-rich motif was identified as the binding site of Roc1, which was confirmed by a functional promoter analysis. Together, Roc1 is a key regulator of cellulose degradation and the first identified in wood-decaying fungi in the phylum Basidiomycota.

## INTRODUCTION

Plants store a considerable amount of energy in lignocellulose, and for that reason wood has been used as a fuel since prehistoric times. Wood is recalcitrant to decay by most organisms, but fungi have evolved ways to degrade part of the lignocellulose into its monomeric constituents. Most wood decay fungi belong to the phylum Basidiomycota, or, more specifically, the class Agaricomycetes. Phylogenetically these are distantly related to the fungi in the phylum Ascomycota such as *Saccharomyces cerevisiae* and *Neurospora crassa*. The last common ancestor of ascomycete and basidiomycete fungi is estimated to have lived over 600 million years ago^1^. Although the Ascomycota harbour potent lignocellulose-degrading fungi, the strongest wood-decaying fungi are found in the Basidiomycota.

Lignocellulose consists of a wide range of components, including cellulose, hemicellulose, pectin, and the aromatic polymer lignin. These polymers are found in the plant cell wall. Fungi can generally easily absorb glucose and other monomers from the growth medium, but lignocellulose requires extensive extracellular enzymatic degradation before the breakdown product can be transported into the cells and metabolized. Wood-degrading fungi have evolved a broad range of hydrolytic enzymes that break down the various components of lignocellulose, including cellulases, hemicellulases, pectinases and oxidative enzymes. Collectively, these plant cell wall degrading enzymes are known as carbohydrate-active enzymes (CAZymes) and are classified into families of Glycoside Hydrolases (GHs), Glycosyl Transferases (GTs), Polysaccharide Lyases (PLs), Carbohydrate Esterases (CEs), and Auxiliary Activities (AAs)^2,3^. A typical genome of a wood-degrading fungus encodes hundreds of CAZymes^4–6^.

Basidiomycete wood decayers can be broadly divided into white rot fungi, which degrade all components of the plant cell wall, and brown rot fungi, which depolymerize cellulose, but leave lignin largely unmodified. However, fungi that show (genotypic and phenotypic) characteristics of both white rot and brown rot fungi have also been identified^4^. Neither white rot fungi nor brown rot fungi form monophyletic groups, and the brown rot lifestyle has evolved several times from white rot fungi^1,4,7^.

Genes that encode CAZymes are generally strictly regulated at the transcriptional level, since their production is energetically expensive and not always needed. The primary mechanism of regulation is carbon catabolite repression (CCR). CCR represses the production of ligninolytic enzymes in the presence of an easily metabolizable carbon source, such as glucose, and ensures that the organism pursues the most energy-efficient mode of growth by ideal resource utilization. CCR is regulated by a highly conserved zinc-finger transcription factor (*CreA*/*Cre1*/*Cre-1*)^8–10^, which functions as a strong inhibitor of gene expression in the presence of simple sugars and has been described in several ascomycetes. The gene is conserved in basidiomycetes^10^ and it indeed plays the same role in the mushroom-forming white rot *Pleurotus ostreatus*^11^.

A wide range of transcription factors act downstream of CCR (i.e. in the absence of simple sugars). Generally, these transcription factors activate gene expression of CAZymes involved in the breakdown of specific polysaccharides. Examples include *xlnR, CLR-1, CLR-2* and *ACE1*. In *Aspergillus*^12^ and *Trichoderma*^13^ the transcription factor *xlnR* regulates (hemi)cellulose degrading enzymes and it has an ortholog in almost all filamentous ascomycetes^14^. In *N. crassa* the transcription factors *CLR-1 and CLR-2* induce the expression of cellulolytic, but not the hemicellulolytic enzymes^15^. In contrast to the aforementioned regulators, *ACE1* is a repressor of cellulolytic and xylanolytic enzyme production in *Trichoderma reesei*^16^. It is important to note that these regulators have only been identified in ascomycete fungi, and that there are no orthologs in basidiomycete fungi^10^. To date, no regulators have been identified in the wood-degrading basidiomycetes that positively regulate CAZymes. In general, very little is known about the regulatory mechanisms involved in plant biomass degradation in the basidiomycetes.

*Schizophyllum commune* is a model system for mushroom-forming fungi in the class Agaricomycetes. Several molecular tools have been developed for this organism, including an efficient gene deletion protocol^17,18^ and a ChIP-Seq protocol for Histone H3 to study the epigenetic landscape^19^. *S. commune* has a wide geographical distribution and is generally found as white fruiting bodies growing on wood. Its mode of wood decay is atypical, since it is not easily classified as either white rot or brown rot^4,6,20^. *S. commune* lacks lignin-degrading peroxidases, limiting its ability to degrade lignin^21^. Still, *S. commune* degrades all wood components, but leaves the middle lamella of the plant cell wall mostly intact^20^.

Here, we describe the identification of a regulator of cellulase expression (Roc1) by comparative genomics and comparative transcriptomics. A deletion strain was unable to efficiently utilize cellulose as a carbon source, and growth was inhibited on other components of wood. Moreover, with ChIP-Seq we identified the binding sites of this transcription factor, which were enriched near CAZymes involved in cellulose degradation. This is the first positive regulator of cellulase expression identified in basidiomycetes.

## MATERIALS AND METHODS

### Strains, media composition and culture conditions

The reference strains used in this study were *Schizophyllum commune* H4-8 (*mat*A43*mat*B41; FGSC 9210) and the compatible isogenic strain H4-8b (*mat*A41*mat*B43) ^21^. Strain *Δku80* was derived from H4-8 and was used for gene deletion^18^. The dikaryotic strains *S. commune* Tattone and *S. commune* Loenen were collected in Tattone (Corsica, France) and Loenen aan de Vecht (The Netherlands), respectively. The monokaryotic strains *S. commune* TattoneD and *S. commune* LoenenD were isolated from these strains by protoplasting. Protoplasting was performed using a *Trichoderma harzianum* Horst lytic enzyme mix as described previously^22^.

The strains were grown at 30°C on a medium comprising (per L): 22 g glucose monohydrate, 1.32 g (NH_4_)_2_SO_4_, 0.5 g MgSO_4_·7H_2_O, 0.12 g thiamine, 1 g K_2_HPO_4_, 0.46 g KH_2_PO_4_, 5 mg FeCl_3._6H_2_O, trace elements and with or without 1.5% agar^23^. For cultures with other carbon sources, glucose was replaced with 1% (w/v) Avicel (cellulose), 1% (w/v) glucose + 1% (w/v) Avicel, 1% (w/v) cellobiose, 1% (w/v) xylan from corncob, 1% (w/v) pectin from apple, 1% (w/v) starch from potato, 2.2% (w/v) xylose, 2.2% (w/v) maltose monohydrate. To improve the visualization of the fungal colonies growing on Avicel, the media was supplemented with 20 µg µl^-1^ Remazol Brilliant Blue R. Liquid cultures were grown in Erlenmeyer flasks at 30°C with shaking at 200 rpm.

For selection on nourseothricin (Bio-Connect, Netherlands), phleomycin (Bio-Connect, Netherlands) or hygromycin (Bio-Connect, Netherlands), the media was supplemented with 15 μg ml^-1^, 25 μg ml^-1^ and 20 μg ml^-1^ antibiotic, respectively.

### Genome sequencing and assembly

To perform genome improvement on assembly version Schco1 of *S. commune* H4-8^21^, the whole genome shotgun assembly was broken down into scaffolds and each scaffold piece was reassembled with phrap and subsequently improved using our Phred/Phrap/Consed pipeline^24,25^. Initially all low-quality regions and gaps were targeted with computationally selected Sanger sequencing reactions completed with 4:1 BigDye terminator: dGTP chemistry (Applied Biosystems). These automated rounds included walking on 3 kb and 8 kb plasmid subclones and fosmid clones using custom primers (400, 3498 and 1183 primers were selected respectively). Following completion of the automated rounds, a trained finisher manually inspected each assembly. Further reactions were then manually selected to improve the genome. Remaining gaps and hairpin structures were resolved by generating small insert shatter libraries of 8 kb-spanning clones. Smaller repeats in the sequence were resolved by transposon-hopping and sequencing 8 kb plasmid clones. 136 fosmid clones were shotgun sequenced and finished to fill large gaps and resolve larger repeats. All these sequencing reactions were generated using Sanger long-read technology.

The genomes of *S. commune* TattoneD and LoenenD were sequenced using 270 bp insert standard fragment Illumina libraries in 2×150 format. The resulting reads were filtered for artifact and process contamination and were subsequently assembled with Velvet^26^. The resulting assembly was used to create *in silico* long mate-pair library with insert 3000 +/- 300 bp, which was then assembled together with the original Illumina library with AllPathsLG release version R42328^27^.

### Gene prediction and functional annotation

The genomes were annotated using the JGI Annotation Pipeline^28,29^, which combines several gene prediction and annotation methods, and integrates the annotated genome into the web-based fungal resource MycoCosm^29^. Before gene prediction, repetitive sequences in the assemblies were masked using RepeatMasker^30^, RepBase library^31^, and the most frequent (>150 times) repeats were recognized by RepeatScout^32^. The following combination of gene predictors was run on the masked assembly: ab initio Fgenesh^33^ and GeneMark^34^; homology-based Fgenesh+^33^ and Genewise^35^ seeded by BLASTx alignments against the NCBI NR database. RNA-Seq data (see below) was used during gene prediction for strains H4-8 and TattoneD, but not for LoenenD. In addition to protein-coding genes, tRNAs were predicted using tRNAscan-SE^36^.

The predicted proteins of the three assemblies were functionally annotated. PFAM version 32 was used to predict conserved protein domains^37^ and these were subsequently mapped to their corresponding gene ontology (GO) terms^38,39^. Secretion signals were predicted with Signalp 4.1^40^ and transmembrane domains were predicted with TMHMM 2.0c^41^. Proteins were considered small secreted proteins when they had a secretion signal, but no transmembrane domain (except in the first 40 amino acids) and were shorter than 300 amino acids. Transcription factors were identified based on the presence of a PFAM domain with DNA binding properties^42^. Proteases were predicted based on the MEROPS database^43^ using a BlastP E-value cutoff of 1e-5. A pipeline based on the SMURF method^44^ was used to predict genes and gene clusters involved in secondary metabolism. SMURF parameter d (maximum intergenic distance in base pairs) was set at 3000 bp. SMURF parameter y (the maximum number of non-secondary metabolism genes upstream or downstream of the backbone gene) was set at 6. CAZymes were annotated with the standalone version of the dbCAN pipeline using HMMdb version 9^45^.

The assemblies of strains TattoneD and LoenenD were aligned to the assembly of H4-8 using PROmer version 3, which is part of the MUMmer package^46^. The setting “mum” was used. Next, a sliding window approach (1 kbp window with 100 bp step size) was taken to determine the percentage of identity across the assemblies of the strains.

### Comparative transcriptomics during growth on various carbon sources

Cultures of strain H4-8 and TattoneD were pre-grown on a Whatman Cyclopore™ polycarbonate (PC) membrane on top of minimal medium containing glucose at 30°C in the dark. After 3 days, the PC membranes containing the cultures were carefully transferred to fresh plates containing solid minimal medium with either glucose, cellulose (avicel) or birchwood (ground to a particle size of 1 to 3 mm) as sole carbon source. After 3 days, the cultures were harvested, lyophilized, powdered in liquid nitrogen, and RNA was extracted using the Zymo Direct-zol RNA MiniPrep kit. The quality of the RNA was assessed with an Agilent 2100 Bioanalyzer. All conditions were analyzed with biological triplicates.

Illumina libraries were generated for RNA-Seq and subsequently sequenced on the Illumina HiSeq-2000 platform in 1×50 bp mode. The exceptions to this were the three glucose replicates from strain H4-8, since these were sequenced in 2×100 bp mode. To avoid any biases during mapping and counting between these samples and the others, only the first 50 bp from the left read pair were extracted from these samples using BBduk (part of the BBMap suite ^47^) and used in the subsequent transcriptome analysis.

The sequence reads were aligned to their respective genome assemblies, *S. commune* H4-8 (version Schco3) or *S. commune* TattoneD (version Schco_TatD_1), using the aligner Hisat version 2.1.0^48^. Default settings were used, with these exceptions: --min-intronlen 20 --max-intronlen 1000. Expression values were calculated as RPKM values (Reads per Kilobase model per Million mapped reads) using Cuffdiff version 2.2.1, which is part of the Cufflinks package^49^. The bias correction method was used while running Cuffdiff^50^. In addition to the cutoff used by Cuffdiff to identify differentially expressed genes, we applied an additional filter of at least a 4-fold change in expression value, as well as at least one condition with an expression value of at least 10 RPKM. Over-representation and under-representation of functional annotation terms in sets of differentially expressed genes were calculated using the Fisher Exact test. The Benjamini-Hochberg correction was used to correct for multiple testing and as a cutoff for significance we used a corrected *p*-value of 0.05.

To compare gene expression, we first identified orthologs between the two strains. Here, proteins are considered orthologs if they show strong homology (having a best bidirectional hit in a blastP analysis applying an E-value cutoff of 1e-10) and if they display syntenic gene order conservation (at least 1 of 4 neighbors should be shared between the strains).

The gene expression profiles were visualized with a heat map generated by the seaborn package for python (https://seaborn.pydata.org). The genes were clustered using the euclidean distance and average linkage methods. The values were scaled for each gene with a z-transformation, resulting in a z-score.

### Conservation of Roc1 in the fungal kingdom

The genome sequences and predicted genes/proteins of 140 previously published fungi were obtained from the publications listed in Table S1. Curating these previously published gene predictions was beyond the scope of this study. Conserved protein domains were identified using PFAM version 32^37^. Roc1 is classified as a putative transcription factor based on the presence of a Zn_2_Cys_6_ DNA binding domain as well as a fungal specific transcription factor domain (Pfam domains PF00172 and PF04082, respectively). We took a multi-step approach to more accurately identify putative Roc1 orthologs. First, we identified transcription factors of the same broad family by selecting proteins with one Pfam domain PF00172 and one Pfam domain PF04082. Next, the sequences of these domains were concatenated and used to reconstruct an initial gene tree of fungal-specific transcription factors. The sequences were aligned with MAFFT 7.310 using auto settings^51^. FastTree 2.1 with default settings was used to calculate the gene tree^52^. Manual inspection of the tree revealed a group of proteins from basidiomycetes and ascomycetes that clustered with Roc1, and these were labeled as candidate orthologs. Next, the full proteins of these candidate orthologs were aligned (instead of only the Pfam domains) with MAFFT and FastTree (as described above). Manual inspection of both the tree and the alignments revealed that in the Agaricomycetes the conserved sequence extended along the entire protein, while in the other basidiomycetes as well as the other phyla conservation was largely restricted to the Pfam domains. For this reason, we constructed a custom HMM model using the full protein alignments of only the Roc1 candidates of the Agaricomycetes. This HMM model was made using HMMER version 3.3.2 (hmmer.org) with default settings. The predicted proteins in the genomes of all 140 fungi were scanned with the HMM, using hmmsearch (part of the HMMER package) with a score cutoff of 500. A new gene tree was calculated as described above, containing the proteins in the previously mentioned tree, as well as any proteins that were additionally identified with the HMM approach. Proteins with this conserved Roc1 HMM domain, Pfam domain PF00172 and Pfam domain PF04082 were considered Roc1 orthologs, while the other candidates were considered distant homologs.

A phylogenetic tree of the 140 species (species tree) was reconstructed using 25 highly conserved proteins identified with BUSCO v2 (dataset ‘fungi_odb9’)^53^. Sequences were aligned with MAFFT 7.307^51^ and well-aligned regions were subsequently identified using Gblocks 0.91b^54^ resulting in 17196 amino acid positions. FastTree 2.1 was used for phylogenetic tree reconstruction using default settings^52^. The phylogenetic species tree was rooted on ‘early-diverging fungi’ (i.e. non-Dikarya). Python toolkit ete3^55^ was used to visualize the gene tree and species trees.

### Deletion of gene *roc1* in strain H4-8

Gene *roc1* (proteinID 2615561 in version Schco3 of the genome of *S. commune*) was deleted using our previously published protocol^17^, which uses pre-assembled Cas9-sgRNA ribonucleoproteins and a repair template to replace the target gene with a selectable marker. The repair template was a plasmid comprising a pUC19 backbone, 1068 bp up-flank of *roc1*, a 1326 bp nourseothricin resistance cassette, 1062 bp down-flank of *roc1*, and a phleomycin resistance cassette. The nourseothricin resistance cassette was cut from plasmid pPV010 using the restriction enzyme EcoRI. The 4112 bp pUC19 backbone and phleomycin resistance cassette were cut from plasmid pRO402 using the restriction enzyme HindIII. The up-flank and down-flank were amplified from genomic DNA of strain H4-8 using primer pairs Roc1UpFw / Roc1UpRev and Roc1DownFw / Roc1DownRev, respectively (Table S2). The 5’ overhangs of these primers were chosen to facilitate Gibson assembly of the four fragments into a single plasmid (NEBuilder HiFi DNA Assembly Master Mix; New England Biolabs). The resulting plasmid pRO405 was verified by restriction analysis and Sanger sequencing.

The sgRNAs were designed on regions downstream and upstream of the up-flank and down-flank of *roc1*, respectively, and one sgRNA was selected for each region as previously described^17^. They were synthesized *in vitro* using the GeneArt Precision sgRNA Synthesis Kit (ThermoFisher Scientific) using *roc1*-specific primers Roc1LeftsgRNAp1 / Roc1LeftsgRNAp2 and Roc1RightsgRNAp1 / Roc1RightsgRNAp2 (Table S2).

Protoplasts of strain *Δku80* were transformed with the pre-assembled ribonucleoproteins and the repair template, as previously described^17^. A first selection was done on minimal medium with 15 μg ml^-1^ nourseothricin. The resistant transformants were transferred to a second selection plate with nourseothricin and subsequently screened on minimal medium with 25 μg ml^-1^ phleomycin. Nourseothricin-resistant transformants that are phleomycin-sensitive are candidates for *roc1* deletion strains, whereas those that are phleomycin-resistant are likely the result of an ectopic integration of the plasmid and therefore undesirable. Six transformants were nourseothricin-resistant, of which four were phleomycin-sensitive. The latter were candidates to have the gene deletion and a confirmation PCR was carried out with primers Roc1-Chk-A and Roc1-Chk-D (Table S2) that amplify the integration locus (resulting in a 5007 bp product in the case of the wild type situation, or a 3532 bp product in the case of a correct gene deletion). One of these *Δroc1Δku80* strains was selected and crossed with the compatible wild type H4-8b in order to eliminate the *Δku80* background. Meiotic basidiospores were collected and the offspring was grown on minimal medium with nourseothricin and 48 out of 72 were resistant. A second selection was done with the 48 nourseothricin-resistant strains on minimal medium with 20 μg ml^-1^ hygromycin and 23 individuals out of 48 were hygromycin-sensitive, indicating that these do not have the *Δku80* background. A nourseothricin-resistant and hygromycin-sensitive strain with the same mating type as H4-8 was selected.

### Complementation of the *Δroc1* deletion

The *roc1* deletion strain was complemented with a plasmid expressing a C-terminally haemagglutinin-tagged version of *roc1*. The promoter and coding sequence of *roc1* were amplified with primers Roc1ChipFw and Roc1ChipRev. Plasmid pPV009 (which contains a C-terminal triple HA tag as well as a phleomycin resistance cassette) was linearized with HindIII. Gibson assembly was used to combine the two fragments into the final complementation plasmid. The protoplasting of *S. commune Δroc1*a and the transformation with the complementation plasmid were carried out as previously described^22^. A first selection was done on minimal medium with 25 μg ml^-1^ phleomycin. Twenty-three transformants were transferred to a second selection plate with phleomycin. Twenty-one of them showed growth and they were initially screened on cellulose (Avicel) plates together with the wild type. Ten transformants had a similar growth when compared to the wild type. A second screening was done on minimal medium and 7 transformants resembled the wild type phenotype. These were subjected to a PCR check with primers Roc1ChipCheckFw and Roc1ChipCheckRev (Table S2). Four transformants showed the desired 4330 bp fragment indicating they might be good candidates for *Δroc1* complementation strains. One of these strains was selected and named *Δroc1 :: roc1-HA*. A western blot was done (see below) and this strain showed a 78 kDa band when grown on cellulose, which was absent in the wild type. This indicated that the Roc1 protein was correctly tagged with the HA-tag.

### Cellulase activity assay

The *S. commune* strains were pre-cultured on a Poretics™ Polycarbonate Track Etched (PCTE) Membrane (GVS, Italy) placed on top of glucose medium for 5 days at 30°C. The mycelia of five cultures for each strain were macerated in 100 ml minimal medium with either 1% cellulose (Avicel) or 5% glycerol (as indicated in the Results section) for 1 minute at low speed in a Waring Commercial Blender. The macerate was evenly distributed to 250 ml Erlenmeyers (20 ml each) containing 80 ml minimal medium with either 1% cellulose (Avicel) or 5% glycerol. Four Erlenmeyers for each strain were placed in an Innova incubator shaker for 10 days at 30°C with shaking at 200 rpm. Samples of the culture medium (1 mL) were collected after 6 days and centrifuged at 9400 g for 10 min. The supernatant was then used for the total cellulase enzyme activity measurement using the filter paper activity (FPase) assay^56^. Total cellulase activity was determined by an enzymatic reaction employing circles with a diameter of 7.0 mm of Whatman No. 1 filter paper and 60 µl of supernatant. The reaction was incubated at 50°C for 72 hours. Next, 120 μL of dinitrosalicylic acid (DNS) was added to the reaction, which was then heated at 95°C for 5 minutes. Finally, 100 µl of each sample was transferred to the wells of a flat-bottom plate and absorbance was read at 540 nm using a BioTek Synergy HTX Microplate Reader. One enzyme unit (FPU) was defined as the amount of enzyme capable of liberating 1 μmol of reducing sugar per minute (as determined by comparison to a glucose standard curve).

### Western blot analysis

A Western blot was performed to confirm that the Roc1 protein was correctly tagged with the haemagglutinin tag. Nine-day old mycelia grown on a Polycarbonate Track Etched (PCTE) Membrane (GVS, Italy) placed on top of 1% Avicel plates were harvested, snap-frozen in liquid nitrogen and ground to a fine powder using a Qiagen Tissue Lyser II (Qiagen, Germany) at 25 Hz for 60 seconds. 120 mg of biomass per sample was boiled in 500 µl of 2x Laemmli sample buffer (4% SDS, 20% glycerol, 10% B-mercaptoethanol, 0.004% bromophenol blue, 0.125M Tris pH 6.8) for 5 minutes, centrifuged for 10 minutes at 10000 g to precipitate cellular debris, and 20 µl of each sample was size separated on a 12% Mini-PROTEAN® TGX Stain-Free™ Precast Gel (Bio-Rad, CA, USA) at 200V for 40 minutes. After electrophoresis the proteins were transferred to a polyvinylidene difluoride membrane (ThermoFisher Scientific, MA, USA) according to the manufacturer’s specification. The membrane was blocked for 1 hour with 5% bovine serum albumin (Sigma-Aldrich, MO, USA) in phosphate buffered saline supplemented with Triton X-100 (PBS-T) (137 mM NaCl, 10 mM Na_2_HPO4, 1.8 mM KH_2_PO4, 2.7 mM KCl, 0.1% Triton X-100) and then incubated for one hour with 1:10000 diluted monoclonal mouse anti-HA antibody (#26183, ThermoFisher Scientific, MA, USA) in PBS-T. After incubation, the membrane was washed five times for 15 minutes with PBS-T. The membrane was incubated for one hour with 1:10000 diluted horseradish peroxidase-coupled goat anti-mouse antibody (#62-6520, ThermoFisher Scientific, MA, USA) in PBS-T and washed again five times for 15 minutes with PBS-T. The antibody binding was imaged with Clarity Western ECL Substrate (Bio-Rad, CA, USA).

### ChIP-seq analysis

Protein-DNA interaction and binding sites of Roc1 were surveyed by chromatin immunoprecipitation followed by next-generation sequencing (ChIP-Seq). The ChIP was performed with Pierce Anti-HA Magnetic Beads (ThermoFisher Scientific, MA, USA) and was adapted from previous studies in human cell lines and *Zymoseptoria tritici*^57,58^ and our recently developed method for ChIP-Seq on Histone H3 (H3K4me2) in *S. commune*^19^. Briefly, monokaryons of strains H4-8 or H4-8 *Δroc1 :: roc1-HA* were grown on medium with Avicel on Poretics™ Polycarbonate Track Etched (PCTE) Membrane (GVS, Italy). After 9 days 10 colonies were collected per replicate and washed twice in Tris-buffered saline (TBS) (50 mM Tris pH 7.5, 150 mM NaCl). The colonies were fixated by vacuum infiltration with 1% formaldehyde in TBS for 10 minutes and the reaction was quenched by vacuum infiltration with 125 mM glycine for 5 minutes. The samples were frozen in liquid nitrogen and homogenized in stainless steel grinding jars in a TissueLyser II (Qiagen) for 2 minutes at 30 Hz. The resulting homogenized mycelium was resuspended in 10 ml cell lysis buffer (20 mM Tris pH 8.0, 85 mM KCl, 0.5% IGEPAL CA-630 (Sigma, MO, USA), 1x cOmplete protease inhibitor cocktail (Roche, Switzerland) and incubated on ice for 10 minutes. The samples were centrifuged at 2500 g for 5 minutes at 4°C and the pellet was resuspended in 3 mL nuclei lysis buffer (10 mM Tris pH 7.5, 1% IGEPAL CA-630, 0.5% sodium deoxycholate, 0.1% SDS, 1x cOmplete protease inhibitor cocktail). The released chromatin was fragmented on ice for 8 minutes with sonication, using a branson sonifier 450 (Emerson, MO, USA) with a microtip at setting 4 with 35% output. To prevent sample degradation, the microtip was cooled for 1 minute every 2 minutes. Pure fragmented chromatin was obtained by collecting the supernatant after centrifugation at 15000 g for 10 minutes at 4°C. As input control, 300 µl of sheared chromatin was stored at -80°C for subsequent DNA isolation (see below). The volume of the remaining sheared chromatin was adjusted to 3 ml with ChIP dilution buffer (0.01% SDS, 1.1% Triton X-100, 1.2 mM EDTA, 16.7 mM Tris pH 8.0,167 mM NaCl, 1x cOmplete protease inhibitor cocktail). The chromatin was immunoprecipitated for 16 hours with 50 µl Anti-HA magnetic beads that were equilibrated with ChIP dilution buffer. After incubation, the beads were collected and subsequently washed in low salt washing buffer (0.1% SDS, 1% Triton, 2 mM EDTA, 20 mM Tris pH 8.0, 150 mM NaCl), twice in high salt washing buffer (low salt washing buffer with 500 mM NaCl), twice in lithium chloride washing buffer (250 mM LiCl, 1% IGEPAL CA-630, 1% sodium deoxycholate, 1 mM EDTA, 10 mM Tris pH 8.0) and twice in TE buffer. During all washing steps the samples were incubated for 5 minutes. After addition of the second lithium chloride washing buffer, the samples were transferred from 4°C to room temperature. The chromatin was eluted from Anti-HA magnetic beads by incubating twice in 250 µl elution buffer (1% SDS, 100 mM NaHCO_3_) for 10 minutes. After ChIP, the input controls were adjusted to 500 µl with water. Both samples and input controls were incubated with 50 µg RNAse A for 1 hour at 50°C and decrosslinked overnight at 65°C with 75 µl reverse crosslinking buffer (250 mM Tris pH 6.5, 62.5 mM EDTA, 1.25 M NaCl, 5 mg ml^-1^ proteinase K (Thermofisher Scientific). DNA was isolated with phenol-chloroform extraction. Briefly, samples were mixed with 1 volume of phenol-chloroform (1:1), samples were centrifuged at 15000 g for 5 minutes and the aqueous phase was collected. This step was repeated 2 times. The extraction was repeated with 1 volume of chloroform, to remove residual phenol. The DNA was coprecipitated with 20 mg glycogen (ThermoFisher, MA, USA) by the addition of 58 µL 3M NaAc pH 5.6 and 1160 µL ethanol and stored at -80°C for 2 hours. The DNA was collected by centrifugation at 15000 g for 45 minutes at 4°C and washed with 1 mL 70% ethanol. Finally, DNA was dissolved in 30 µL TE buffer. Next, the DNA was purified with the ChargeSwitch gDNA Plant Kit (Thermofisher Scientific, MA, USA) according to manufacturer’s specifications and eluted in 50 µl ChargeSwith Elution Buffer. The DNA samples were amplified and barcoded with the NEXTFLEX Rapid DNA-Seq library kit (Bioo Scientific, TX, USA) according to manufacturer’s specifications without size selection. The DNA concentration was determined with the NEBNext Library Quant Kit for Illumina (New England Biolabs, MA, USA) and pooled in equimolar ratios with unique barcodes for each sample. The libraries were sequenced on an Illumina NextSeq 500 with paired-end mid output of 75 bp by the Utrecht Sequencing Facility (USeq, www.useq.nl).

The paired-end reads of the controls and samples were aligned to the *S. commune* H4-8 reference genome (version Schco3^21^) with bowtie2 (version 2.3.4.1)^59^. Reads with multiple alignments and a quality score < 2 were removed with samtools (version 1.7) ^60^. Optical duplicates were flagged with picard tools (http://broadinstitute.github.io/picard/) (version 2.21.6) and removed with samtools. Peaks in both WT H4-8 and H4-8 *Δroc1::roc1-HA* were identified with macs2 (version 2.2.3)^61^. To filter out any non-specific binding, peaks identified in both the WT and *Δroc1 :: roc1-HA* strains were excluded from the analysis. Peaks that overlapped repetitive regions (including transposons) were removed. The peaks were associated with a gene if they were within 1000 bp of the predicted translation start site. The correlation between replicates was determined with the R package DiffBind (version 2.12.0).

### Motif discovery

STREME (which is part of the MEME Suite^62^) was used to identify conserved motifs in the ChIP-Seq peaks. STREME looks for ungapped motifs that are relatively enriched in a set of sequences compared to negative control sequences. The 200 bp region around the center of the peak was analyzed for enriched motifs, with 10000 regions of the same length from across the genome as negative sequence set. The minimum motif length was set to 5 bp and the maximum motif length to 25 bp. The location of these motifs in the ChIP-Seq peaks was determined with FIMO (which is part of the MEME Suite^62^). The GC content along the ChIP-Seq peaks was determined with a sliding 25 bp window (step size 5 bp) and averaging the GC content for that window in all ChIP-Seq peaks. To plot the location of the conserved motifs, the peaks were first divided into bins of 20 bp. Next, the density of the motifs along the ChIP-Seq peaks was determined for each bin by dividing the number of motifs in that bin (in all ChIP-Seq peaks) by the total number of ChIP-Seq peaks.

### Functional promoter analysis

Several lengths (approximately 100 bp, 200 bp, 300 bp and 700 bp) of the promoter (defined here as the region located 5’ of the predicted translation start site) of gene *lpmA* (proteinID 1190128) were amplified from H4-8 genomic DNA with the primers GbGH61Pr700Fw (or the primer corresponding to the length) and GbGH61PrRev. The gene encoding the red fluorescent protein dTomato^63^ was amplified from plasmid pRO151^64^ with primers GbdTomatoFw and GbdTomatoRev. The primers used are listed in Table S2. The plasmid backbone (which comprises a nourseothricin resistance cassette) was cut from vector PTUB750_SS3_HC_iT3_Nour_p20^65^ with restriction enzymes HindIII and BamHI. The components of the construct were joined using the Gibson Assembly method (NEBuilder HiFi DNA Assembly Master Mix, New England Biolabs, MA, USA), resulting in the *dTomato* gene under the control of several lengths of the promoter of *lpmA* (plasmids pRO311, pRO312, pRO313 and pRO305, which contain the promoter length of 100 bp, 200 bp, 300 bp and 700 bp, respectively). A motif in the 300 bp promoter (in plasmid pRO313) was changed from CGGACCG to ATTAAAT by site directed mutagenesis using Gibson Assembly (NEBuilder HiFi DNA Assembly Master Mix) and primers GbGH61Pr300MutFw and GbGH61Pr300MutRv (Table S2), resulting in plasmid pPV049. Protoplasts of strain H4-8b were transformed with these dTomato reporter constructs, as previously described^22^. Nourseothricin-resistant transformants were selected for further analysis under the fluorescence microscope.

### Fluorescence microscopy and sample preparation

Mycelia were grown in triplicate for 72h at 30°C on a Poretics™ Polycarbonate Track Etched (PCTE) Membrane (GVS, Italy) placed on top of solid minimal medium. Microscopy samples were prepared by carefully scraping mycelium of the colony from the PCTE membrane with a scalpel and placing it on a slide with a drop of water for adherence and a cover slip. The dTomato fluorescence was detected with an Axioskop 2 plus microscope (Zeiss, Germany) equipped with a 100 Watt HBO mercury lamp and a sCMEX-20 Microscope Camera (5440×3648 pixels) using the TRITC (Tetramethyl Rhodamine Iso-Thiocyanate) filter (excitation at 550 nm and emission at 573nm). The images were taken using ImageFocus Alpha software (24-bit color depth).

## RESULTS

### Comparison of growth profile of three strains of *S. commune* on various carbon sources

The growth profile of *Schizophyllum commune* was determined on carbon sources associated with wood (including cellulose, hemicellulose, and pectin) and other carbon sources (including glucose, maltose and starch) (Figure 1). The reference strain H4-8 was compared to strains LoenenD and TattoneD. The latter two are haploid (monokaryotic) strains that were obtained (by protoplasting) from the dikaryotic wild isolate strains collected in Loenen aan de Vecht (Netherlands) and Tattone (Corsica, France). Strain LoenenD displayed reduced growth on maltose, starch, xylose, xylan and cellulose, but improved growth on pectin and cellobiose compared to H4-8 (Figure 1). In contrast, the growth profile of strain TattoneD was more similar to that of strain H4-8, with the notable exceptions of cellulose (TattoneD grew slower than H4-8) and pectin (TattoneD grew faster than H4-8). Together, there is considerable phenotypic diversity between the various strains of *S. commune*.

**Figure 1:**
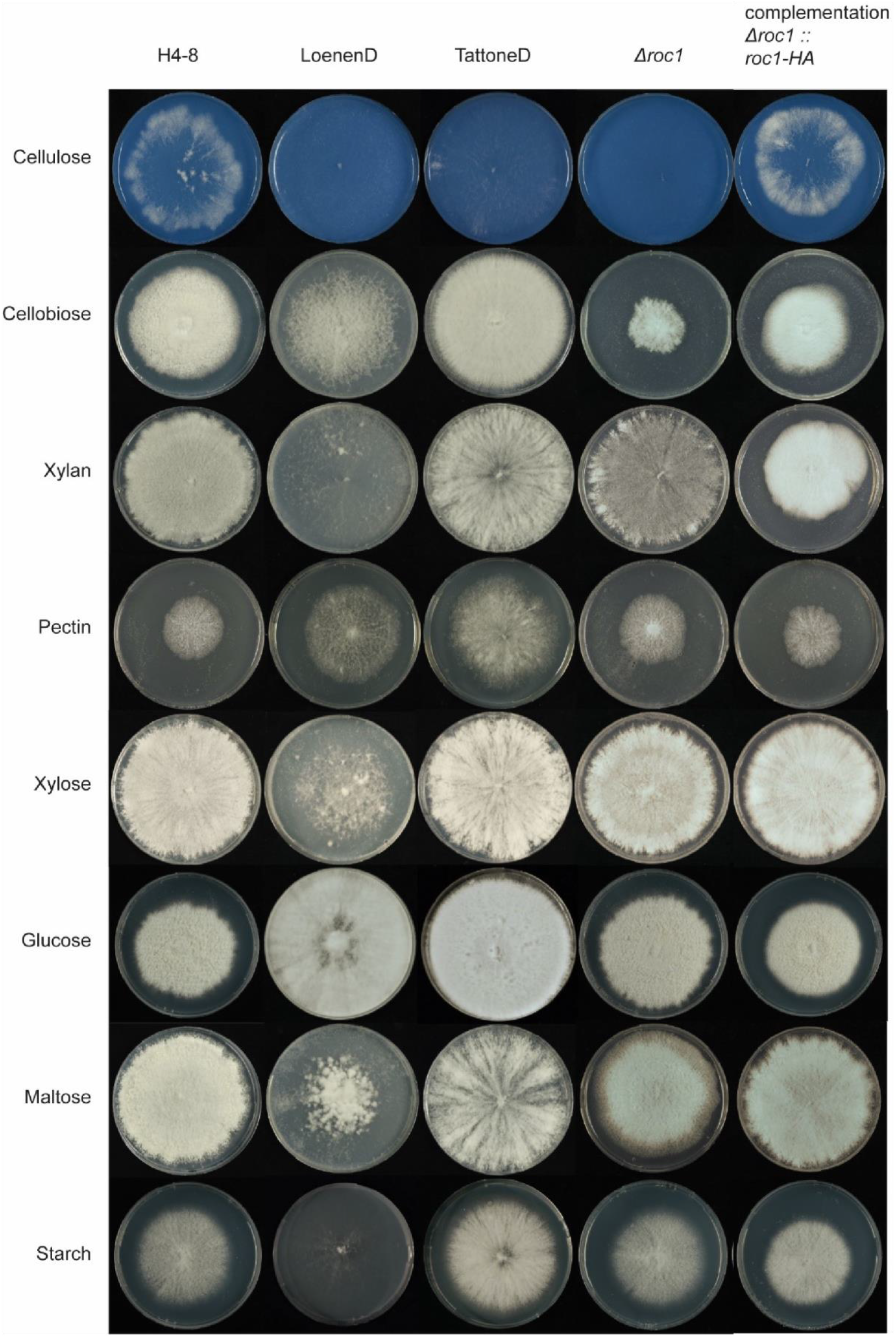
Growth phenotype of *S. commune* strains on various carbon sources. Reference strain H4-8 and wild isolate strains LoenenD and TattoneD displayed high phenotypic plasticity regarding growth on these carbon sources. Strain LoenenD showed reduced growth on maltose, starch, xylose, xylan and cellulose (Avicel), but improved growth on pectin and cellobiose compared to the reference strain H4-8. In contrast, the growth profile of strain TattoneD was more similar to that of strain H4-8, with the notable exceptions of cellulose (TattoneD grew slower than H4-8) and pectin (TattoneD grew faster than H4-8). Deletion strain Δ*roc1* showed strongly reduced growth on cellulose and cellobiose, when compared to its parent strain H4-8. This phenotype was rescued when the deletion was complemented. All strains were grown from a point inoculum for 7 days (glucose) and 11 days (other carbon sources) at 30°C. The cellulose medium was stained blue with Remazol Brilliant Blue R to enhance the visibility of the white mycelium on the white cellulose medium (the dye did not affect growth; data not shown).

### Genome sequences of three strains of *S. commune*

The genome sequence and annotation of strain H4-8 were previously published^21^ and we here report an updated version (Schco3). Moreover, we sequenced strains TattoneD and LoenenD and generated draft assemblies and annotations (Table S3).

The original Sanger-sequenced assembly of strain H4-8 (version Schco1) was improved by extensive targeted gap-sequencing and manual reassembly. This reduced the number of scaffolds from 36 to 25, and considerably reduced the percentage of assembly gaps from 1.43% to 0.15%.

Furthermore, a new set of gene predictions was generated using the RNA-Seq data used for the comparative transcriptomics described below. This raised the gene count from 13181 to 16204. The coding content of the assembly (i.e. the percentage of the assembly consisting of coding sequence) increased from 45.81% to 52.89%, indicating that genes that were missed in the original annotation were added to the new set. All statistics regarding the functional annotation of the predicted genes improved in the new annotation, indicating that the new gene set is more complete (i.e. more predicted genes were assigned a functional annotation) (Table S3). The BUSCO completeness score improved to 99.13%, further showing that the new assembly and gene predictions are of high quality. This new assembly and set of gene predictions will therefore be a valuable tool for functional analysis of this important model system of mushroom-forming fungi.

Draft assemblies and gene predictions were generated for strains TattoneD and LoenenD. Although both are more fragmented than the assembly of reference strain H4-8, the corresponding sets of gene predictions are similarly complete, as determined by BUSCO (98.62% and 99.31%, respectively). Illumina-sequenced genomes are generally more fragmented than Sanger-sequenced genomes, especially regarding repeat-rich regions^66^. This is reflected in the lower percentages of repetitive content for strains TattoneD and LoenenD, compared to H4-8. Importantly, the coding content of the assemblies are in a similar range, indicating that the set of gene predictions is reliable.

The three strains displayed a high degree of sequence diversity at the level of the genome (Figure 2A and 2B). Large parts of the genome display less than 95 % similarity (over a 1 kb sliding window). In some cases (e.g. scaffolds 12, 15 and 19) the (sub)telomeric regions of strain H4-8 are not found in strains TattoneD or LoenenD. Despite this high degree of sequence diversity among the three strains, the majority of genes are conserved between the strains (Figure S1). The set of predicted carbohydrate-active enzymes (CAZymes) is remarkably similar between the strains (Figure 2C and Table S4). Therefore, the difference in growth profile on the various carbon sources cannot be easily explained by the CAZyme gene counts.

**Figure 2.**
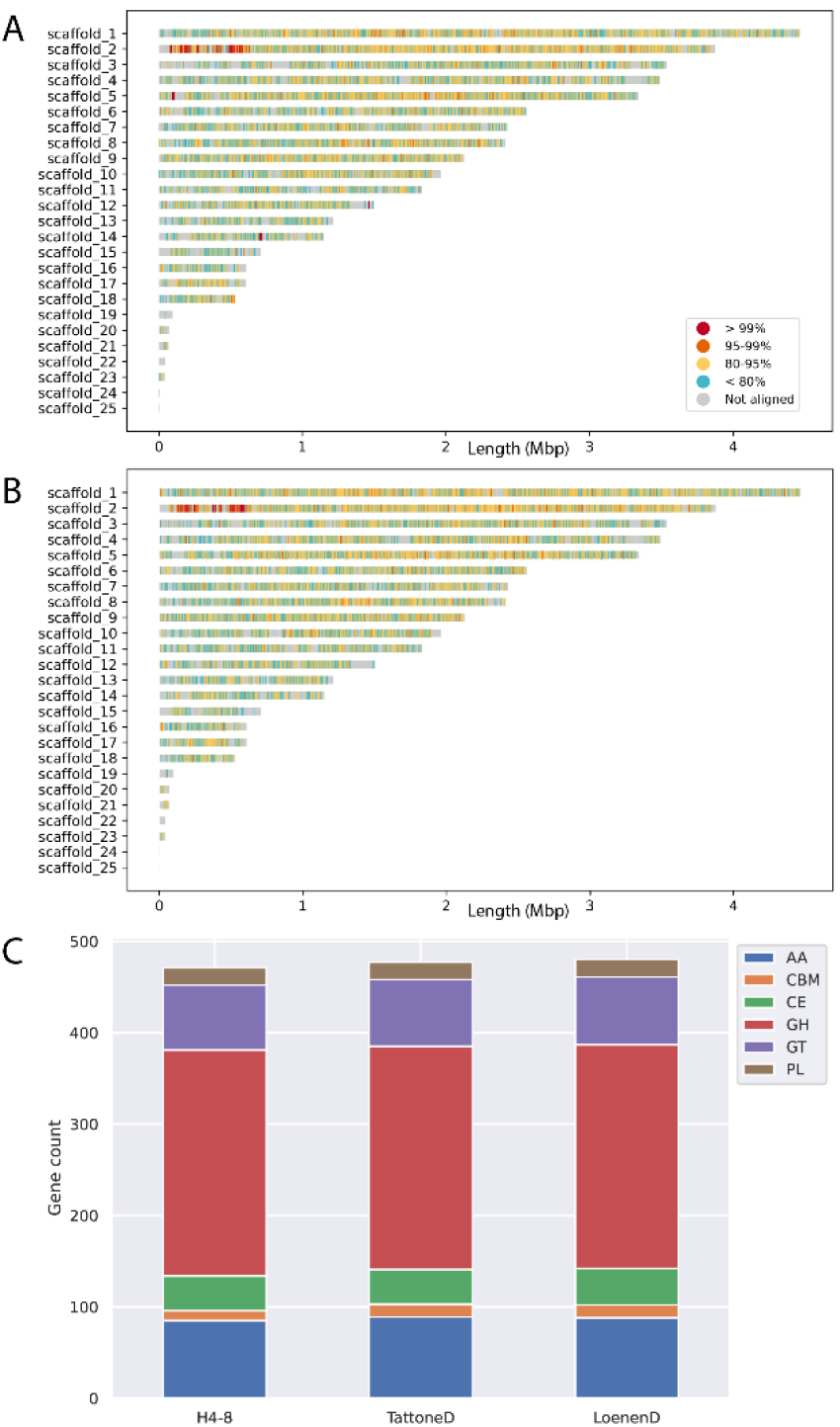
Conservation between the reference assembly of strain H4-8 and the assemblies of strains (**A**) TattoneD and (**B**) LoenenD. Even though these are strains of the same species, their assemblies display a high degree of variation. **C**. The number of predicted genes involved with plant cell wall degradation is very similar between the strains. These CAZymes are classified in subfamilies. GH: Glycoside Hydrolases; GT: Glycosyl Transferases; PL: Polysaccharide Lyases; CE: Carbohydrate Esterases; AA: Auxiliary Activities; CBM: carbohydrate-binding modules

### Comparative transcriptomics reveals conserved expression responses to lignocellulosic carbon sources

We performed a comparative transcriptomics analysis to determine whether, despite the high level of phenotypic and sequence diversity, there is a conserved expression response to lignocellulosic carbon sources. Strains H4-8 and TattoneD were pre-grown on minimal medium with glucose as carbon source and after 3 days the colonies were transferred to medium containing either glucose, cellulose (Avicel), or wood. After 3 days of exposure to this carbon source the colonies were harvested, RNA was isolated, and RNA-Seq was performed (Table S5). The heat maps of the expression profiles are depicted in Figure S2.

Glucose does not require extracellular breakdown by CAZymes, in contrast to the polymeric compounds cellulose and wood. Therefore, the most relevant differences in expression profiles were expected between samples grown on glucose and either cellulose or wood. Indeed, 166 and 210 genes of strains H4-8 and TattoneD were up-regulated when growth on cellulose when compared to glucose, respectively. Similarly, 468 and 500 genes of these strains were up-regulated on wood, respectively (Figure S2).

Next, we performed a comparative transcriptomics analysis on the two strains on various carbon sources, in an effort to identify conserved responses. Orthologs were identified between the two strains and their regulation profile was compared (Figure 3). Again, we focused the analysis on expression on cellulose when compared to glucose (Figure 3A) and on wood when compared to glucose (Figure 3B). Orthologs in the upper right corner of Figures 3A and 3B displayed a conserved expression profile on the corresponding carbon sources and many of those are predicted CAZymes. The orthologs annotated as CAZymes showed a more conserved response between the strains (Pearson correlation of 0.88 and 0.88, on cellulose and wood, respectively) than for the full set of genes (Pearson correlation of 0.54 and 0.66, on cellulose and wood, respectively). The expression profile of transcription factors was less conserved between the strains than the CAZymes (Pearson correlation of 0.5 and 0.7, on cellulose and wood, respectively). In fact, only one transcription factor displayed a conserved expression profile in both strains on cellulose and wood.

**Figure 3.**
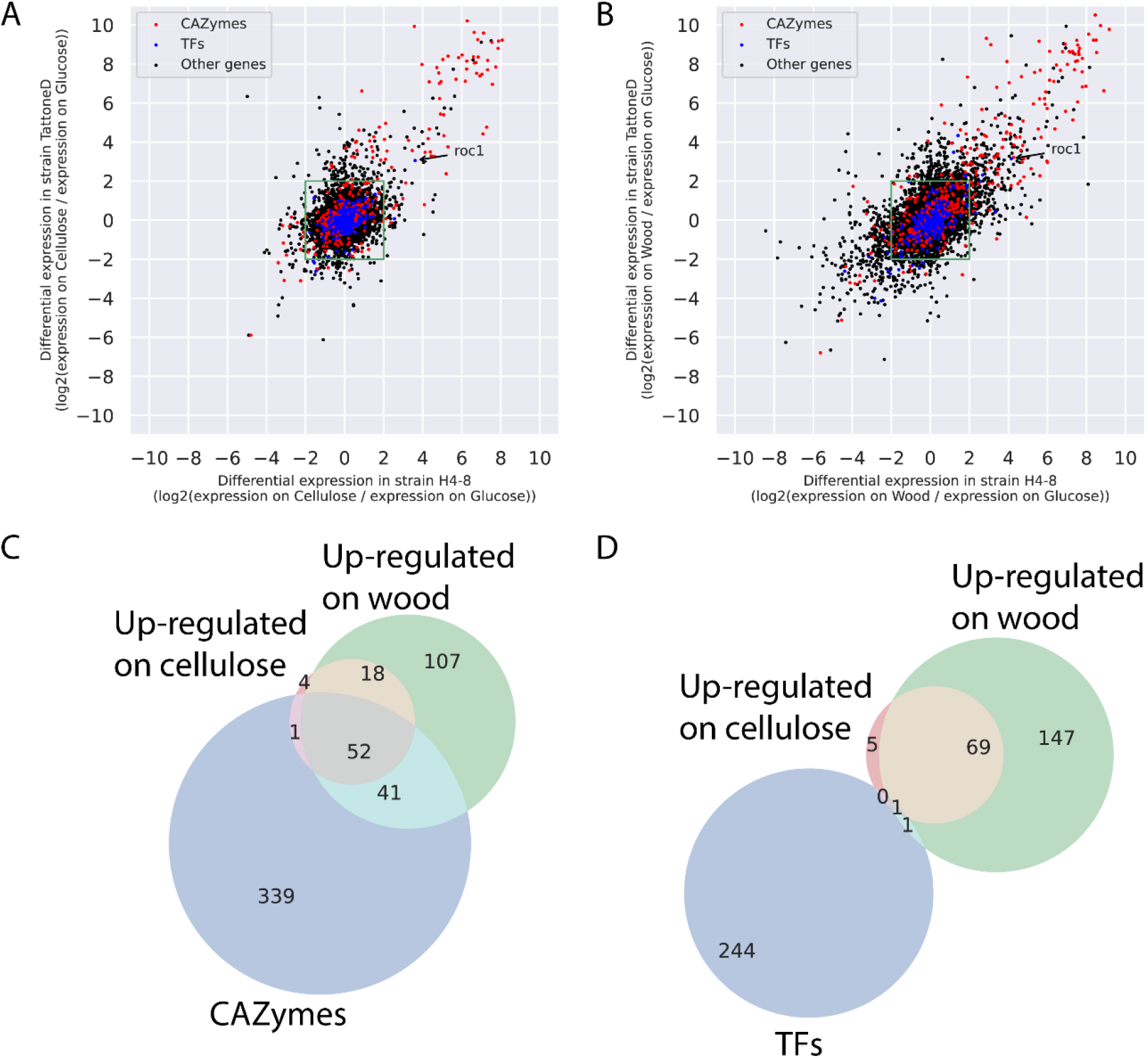
Comparative transcriptomics in strains H4-8 and TattoneD. **A**. Expression of orthologs in the two strains when grown on cellulose, compared to glucose. Orthologs in the green box are not differentially expressed in either strain. Orthologs in the top right quadrant are up-regulated on cellulose in both strains, indicating that they show a conserved response. Many of these orthologs are CAZymes, and only one ortholog is a transcription factor (*roc1*). In general, the response of CAZymes is more conserved than that of other genes **B**. As in (A), but expression on wood when compared to glucose. **C**. VENN diagram of orthologs that are annotated as a CAZyme, are up-regulated on cellulose in both strains, or are up-regulated on wood in both strains (when compared to on glucose). Orthologs that are up-regulated on cellulose in both strains are largely a subset of orthologs up-regulated on wood. Moreover, a considerable number of the up-regulated orthologs are annotated as CAZyme. **D**. As in (C), but for orthologs annotated as transcription factors. Only one transcription factor (*roc1*) was up-regulated in both strains on both cellulose and wood.

Orthologs that are up-regulated on complex carbon sources in one strain, but not in the other strain, (i.e., the dots above and to the right of the green square in Figures 3A and 3B) do not show a conserved expression response. Therefore, those genes may explain part of the difference in phenotype displayed by these strains on complex carbon sources. Furthermore, orthologs that are up-regulated in both strains during growth on cellulose or wood but that do not have a CAZyme annotation (i.e., the black dots in the upper right corners of Figure 3A and 3B) may represent novel CAZymes, or other genes involved in growth on complex carbon sources.

Next, we focused on the orthologs that displayed a conserved expression response (i.e., in both strains) to complex carbon sources when compared to growth on glucose (Figures 3C and 3D). Orthologs that were up-regulated on cellulose in both strains were largely a subset of the orthologs that were up-regulated on wood, and a considerable number of those were CAZymes (Figure 3C). This indicates that the expression program that is activated during growth on cellulose is also activated during growth on wood. However, on wood a large number of additional genes were also up-regulated and were likely involved in the degradation of the complex set of polymers present in this substrate.

Transcription factors, on the other hand, were not found to a large extent in the conserved changes in gene expression (Figure 3D). In fact, only one transcription factor was up-regulated on both cellulose and on wood (when compared to on glucose) in both strain H4-8 and TattoneD (protein ID Schco3|2615561 and Schco_TatD_1|232687; Figure 3A, 3B and 3D). In strain H4-8 the expression is up-regulated 13-fold and 18-fold on cellulose and wood, respectively, when compared to glucose (Table S5). We hypothesized that this transcription factor (from here on named Roc1 for ‘regulator of cellulases’) is involved in the regulation of gene expression during growth on lignocellulose.

### Regulator Roc1 is only conserved in the class Agaricomycetes

Roc1 is classified as a putative transcription factor based on the presence of a Zn_2_Cys_6_ DNA binding domain and a fungal-specific transcription factor domain (Pfam domains PF00172 and PF04082, respectively). These domains are frequently found together and in most fungi this is the most common family of transcription factors^10^ with many (distant and functionally unrelated) members across the fungal kingdom. Examples include GAL4 in *S. cerevisiae*^67^ and XlnR in *Aspergillus niger*^68^. The reference genome of *S. commune* (strain H4-8) encodes 41 members of this transcription factor family.

The genomes of 140 fungi from across the fungal tree were analyzed for orthologs of Roc1 (Table S1). Since the fungal-specific Zn_2_Cys_6_ transcription factor family is large, numerous homologs of Roc1 are found in each genome. We distinguished between homologs and orthologs using a gene tree-based approach, combined with the location of conserved domains (Figure S3). Roc1 orthologs were only found in members of the class Agaricomycetes in the phylum Basidiomycota (Figure S4). These orthologs clustered closely together in the gene tree (having short branch lengths) and were highly conserved across the length of the protein (Figure S3).

### A Δ*roc1* strain is incapable of efficiently utilizing cellulose as a carbon source

A Δ*roc1* strain was generated in strain H4-8 using our recently published CRISPR/Cas9 genome editing protocol^17^ by replacing 2759 bp (which includes the *roc1* coding sequence) with a nourseothricin resistance cassette.

Growth of Δ*roc1* on cellulose (Avicel) was strongly reduced when compared to the reference (Figure 1). Only a very thin mycelium was observed and no aerial hyphae were formed. Moreover, growth on cellobiose and xylan was also reduced, although to a lesser extent than on cellulose. Both these carbons sources are found in lignocellulose. In contrast, Δ*roc1* displays no phenotype when grown on glucose, maltose, starch, pectin and xylose when compared to the reference H4-8.

The wild type phenotype was largely rescued when the Δ*roc1* strain was complemented with the *roc1* coding sequence under control of its own promoter (Figure 1), confirming that the phenotype was caused by the deletion of *roc1*. The coding sequence included a C-terminal haemagglutinin tag, allowing us to use the complementation strain for ChIP-Seq with a commercially available anti-HA antibody (see below).

### Total cellulase enzyme activity in the growth medium is strongly reduced in Δ*roc1*

We assessed whether the strongly reduced growth of the *Δroc1* strain on cellulose coincided with reduced cellulase enzyme activity in the growth medium. Biomass was pre-grown in liquid minimal medium with glucose, washed and subsequently used to inoculate liquid shaking cultures containing cellulose as carbon source. After 6 days, the cellulase activity in the growth medium was determined (Figure 4). Compared to the reference strain H4-8, the *Δroc1* strain had strongly reduced cellulase activity in the growth medium. Moreover, this phenotype was rescued in the complemented Δ*roc1* strain. The lack of growth of *Δroc1* on cellulose (Figure 1) can be explained by the low cellulase activity in this strain, since these cellulases are required to break down cellulose. Furthermore, it suggests that Roc1 regulates the expression of cellulose-degrading genes.

**Figure 4.**
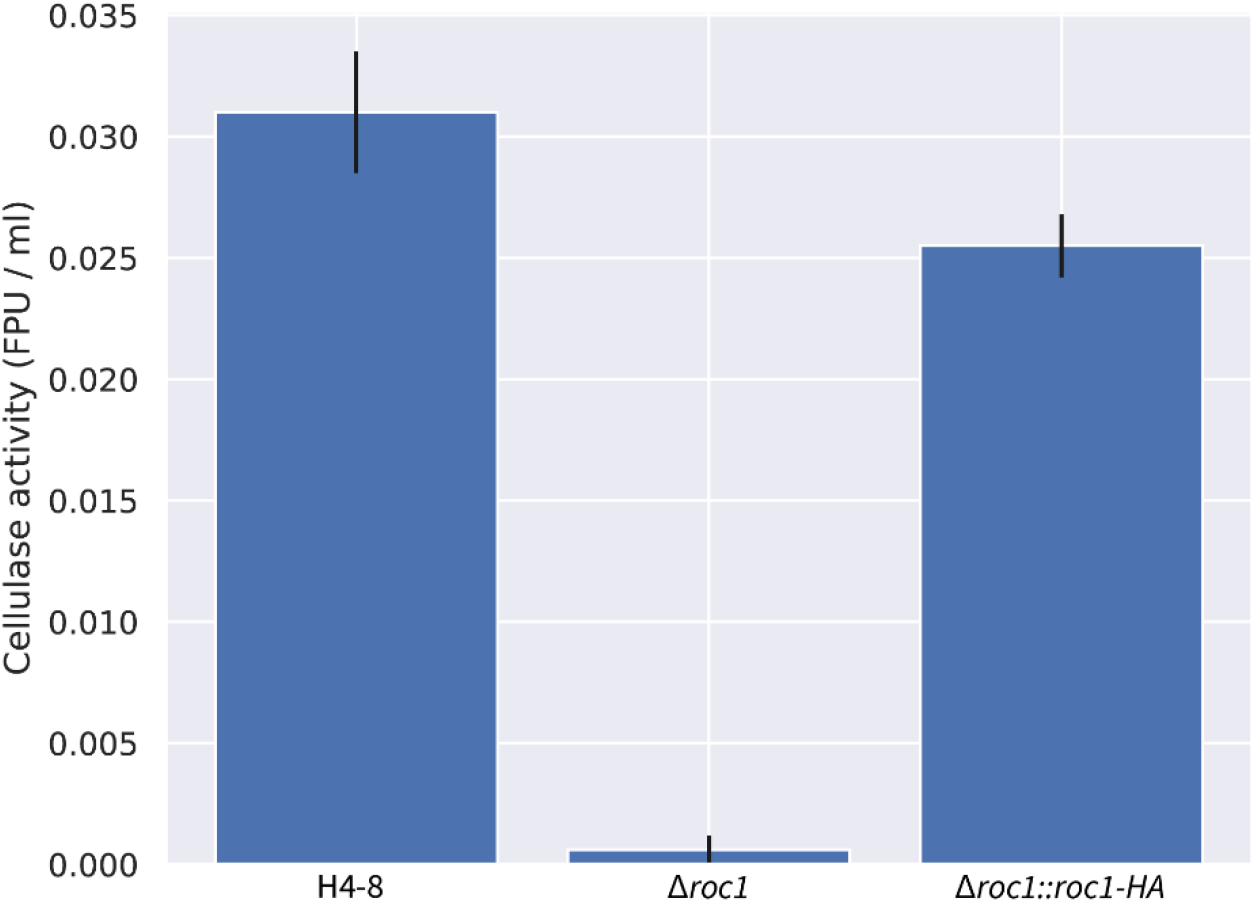
Total cellulase activity of *S. commune* strains in cellulose liquid shaking cultures. There is almost no activity in the Δ*roc1* strain when compared to the reference H4-8. This phenotype is largely rescued upon complementation of the gene. All cultures were pre-grown on glucose medium to ensure that sufficient biomass was present, transferred to cellulose medium, and grown for 6 additional days.

### ChIP-Seq reveals binding sites of Roc1 in promoters of cellulases

Transcription factors may regulate the expression of genes in a direct (e.g., by binding to their promoter) or indirect manner (e.g., by regulating other transcription factors that in turn directly regulate these genes). A ChIP-Seq analysis was performed to identify the binding sites of Roc1 in the genome, allowing us to determine whether Roc1 directly binds the promoters of cellulase genes.

The Roc1 transcription factor was tagged with a haemagglutinin (HA) tag and expressed in the deletion strain (resulting in strain *Δroc1* :: *roc1-HA*), allowing the ChIP-Seq to be performed using commercially available antibodies against the HA-tag (Figure S5). Since this tagged version can complement the phenotype of the *roc1* deletion (Figures 1 and 4), it can be concluded that the HA-tag does not interfere with the function of Roc1.

The strains H4-8 and *Δroc1* :: *roc1-HA* were grown on medium containing cellulose, and the chromatin immunoprecipitation (ChIP) procedure was performed to isolate the DNA to which Roc1 binds. This DNA was subsequently purified and sequenced using Illumina technology. The resulting sequence reads were aligned to the assembly of strain H4 -8 and peaks were identified, which may be considered binding sites of Roc1.

A total of 1474 binding sites of Roc1 were identified during growth on cellulose, which were associated with 1125 unique genes (Table S6). CAZyme genes as a group were not enriched among those genes (p > 0.05), but, in contrast, specific CAZyme families were strongly enriched (Table S7), as well as genes that were up-regulated on cellulose (Figure 5A). A notable family of genes with binding sites of Roc1 are the lytic polysaccharide monooxygenases (LPMOs), also annotated as CAZyme family AA9 (auxiliary activity 9). This family was previously shown to be involved in cellulose degradation^69^. Of the 22 LPMO genes encoded in the genome, 12 were associated with a Roc1 binding site. Moreover, 10 of these 12 were up-regulated during growth on cellulose when compared to growth on glucose. This indicates that Roc1 directly binds to the promoters of these genes during growth on cellulose, activating their expression. The GH3 and GH5 CAZyme families were also over-represented among the genes with a Roc1 binding site (Table S7). Both these glycoside hydrolase families comprise a diverse group of enzyme activities, several of which are involved in (hemi)cellulose degradation. GH3 includes members with reported β-glucosidase (involved in cellulose degradation) and xylan 1,4-β-xylosidase (involved in hemicellulose degradation), while GH5 includes members with reported endo-β-1,4-glucanase (involved in cellulose degradation)^3,70^. Combined, these activities may explain why the Δ*roc1* strain can no longer utilize cellulose as a carbon source and displays slower growth on hemicellulose.

**Figure 5.**
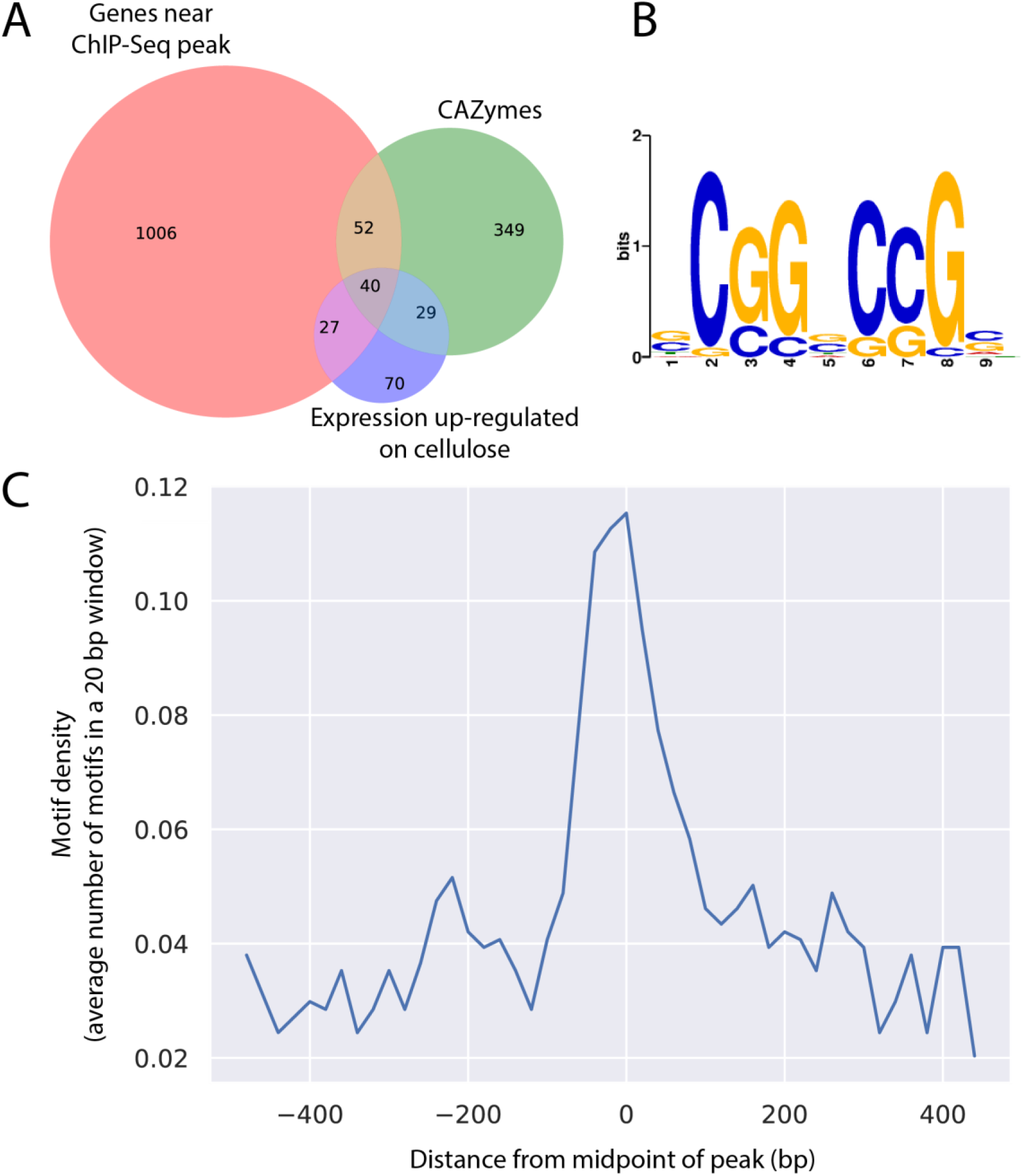
**A**. Venn diagram depicting the overlap between the sets of genes that are associated with a Roc1 binding site, CAZymes, and genes that are up-regulated on cellulose (when compared to glucose). **B**. Conserved motif identified in the binding sites of Roc1. **C**. The binding site in (B) is enriched in the center of the ChIP-Seq peaks.

Remarkably, gene *roc1* itself is also associated with a Roc1 binding site (Table S6). This indicates that Roc1 regulates its own expression, perhaps in a positive feedback loop. Moreover, transcription factor genes in general are enriched among genes associated with a Roc1 binding site. Of the 296 transcription factor genes encoded in the genome, 52 had a nearby Roc1 binding site (Table S7). Few of these 52 transcription factors have been described previously, with the exception of C2h2 and Gat1, which play a role in various aspects of mushroom development ^71^. It should be noted that these two genes were not up-regulated on cellulose or wood, when compared to growth on glucose.

### Conserved GC-rich motif in the Roc1 ChIP-Seq peaks

The peaks identified in the ChIP-Seq analysis were analyzed to identify a conserved motif that represents the binding site of Roc1. The GC content of the 500 bp sequence around the top of each Roc1 peak was determined (Figure S6). There was a marked increase in GC content near the middle of the peaks, which indicates that the Roc1 binding site is GC-rich. Based on the GC curve, we further limited the search to 200 bp around the top of each Roc1 peak. The sequences were analyzed with STREME, which attempts to find conserved motifs that are over-represented in the peak sequences, compared to a representative negative set (i.e., other genomic sequences). This led to the identification of a GC-rich motif (Figure 5B) that was present in 989 of the 1427 peaks and significantly enriched compared to the negative control sequences (p = 2.8·10^−8^ in a Fisher’s exact test). Furthermore, this motif was found most frequently in the center of the identified peak (Figure 5C), as would be expected for the binding site.

### Functional promoter analysis of a CAZyme of the AA9 family reveals the Roc1 binding site

Twelve of the 22 members of the lytic polysaccharide monooxygenase family (LPMOs; CAZyme family AA9) were up-regulated during growth on both cellulose and wood when compared to glucose. Moreover, Roc1 had direct binding sites in the promoters of 12 of the 22 LPMO genes, as determined by ChIP-Seq. Ten of the 22 LPMO genes were both up-regulated on cellulose and had a Roc1 ChIP-Seq peak in their promoter, which shows that there is a strong correlation between these. One of these genes, *lpmA* (protein ID 1190128), was strongly up-regulated on cellulose and wood, compared to growth on glucose (246 and 206-fold, respectively; Table S5). The peak from the Roc1 ChIP-Seq was located upstream of the translation start site (Figure 6A).

**Figure 6.**
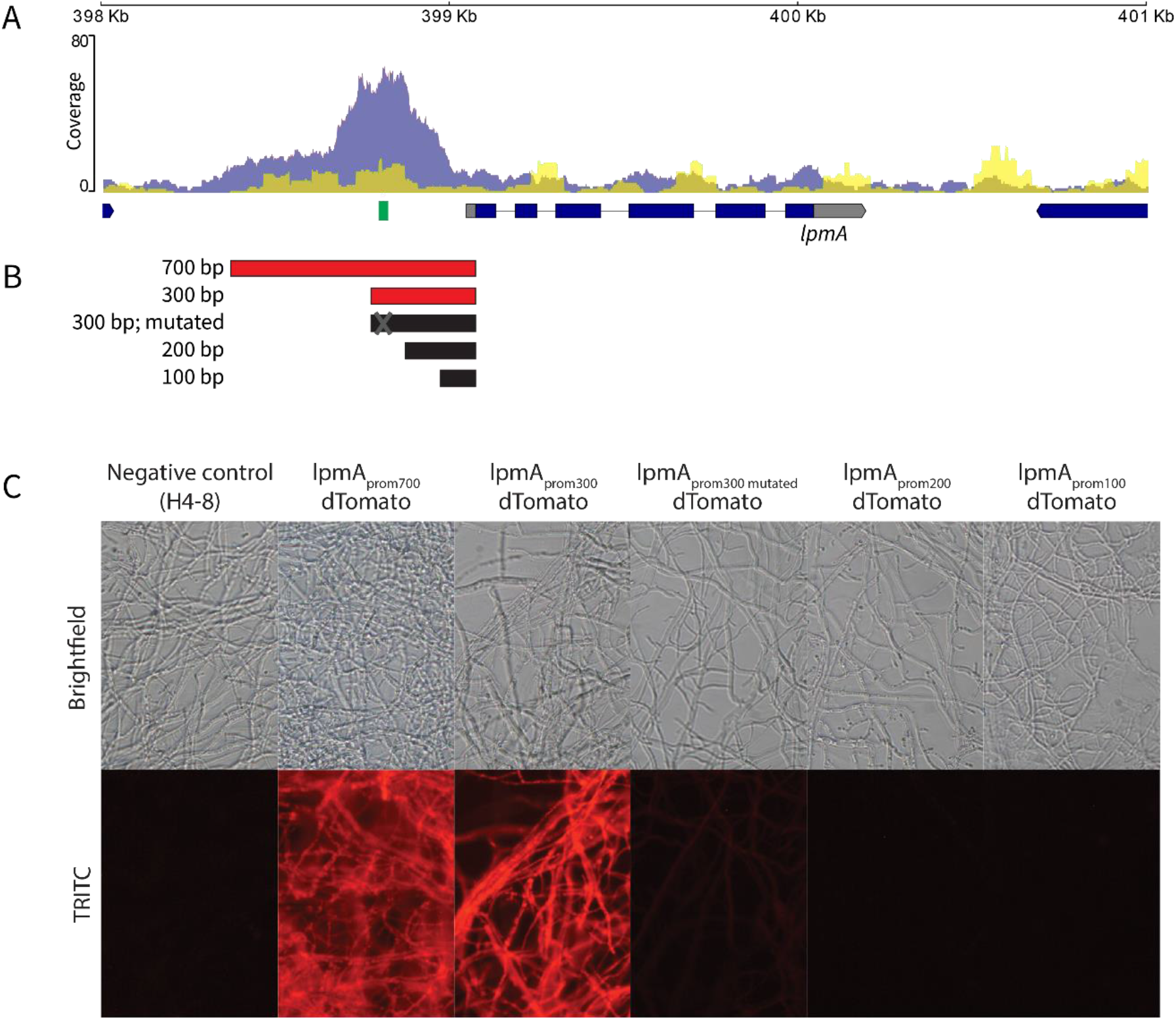
Functional promoter analysis of the lytic polysaccharide monooxygenase *lpmA* gene (protein ID 1190128). **A**. ChIP-Seq read coverage curve in the locus of *lpmA*. The blue curve represents the coverage of the Roc1 ChIP-Seq reads, while the yellow curve represents the negative control. There is a peak in the promoter region up-stream of the *lpmA* coding sequence. The location of the conserved motif (representing the Roc1 binding site; Figure 5B) is indicated in green. **B**. Five regions in the promoter of *lpmA* (5’ of the coding sequence) were tested for their ability to drive expression and fluorescence of dTomato. Active promoter fragments are indicated in red, and inactive promoter fragments in black. **C**. Reporter strains with the dTomato gene under control of the promoters in (B), grown on cellulose. The 700 and 300 bp promoters can drive dTomato expression and fluorescence, but the 200 and 100 bp promoters cannot. The 300 bp promoter in which the Roc1 binding motif had been mutated was not able to drive dTomato expression and fluorescence to the same extent, since only weak fluorescence is observed. When grown on glucose, no fluorescence was observed in any of these strains (Figure S7).

We performed a functional promoter analysis to further investigate the expression dynamics and to locate the transcription factor binding sites. The promoter of *lpmA* was used to drive expression of the red fluorescent reporter protein dTomato (Figure 6B). The 700 bp promoter (i.e., the 700 bp upstream of the predicted translation start site) could drive expression of dTomato when growing on cellulose, resulting in red fluorescent mycelium (strain *lpmA*_prom700_*-dTomato;* Figures 6C). No fluorescence was observed when grown on glucose or a combination of glucose and cellulose (Figure S7). Next, we produced similar reporter constructs with promoter lengths of 300, 200 and 100 bp. The promoter of 300 bp could still drive dTomato expression on cellulose, but, in contrast, the promoters of 200 bp and 100 bp could not (Figures 6B and 6C). This indicates that the region between 300 and 200 bp upstream of the translation start site contains an important regulatory element. This region corresponds with the peak in the ChIP-Seq data and therefore a binding site of Roc1. Moreover, the GC-rich conserved motif (Figure 5B) is located in this region (Figure 6A). This motif is also conserved in the promoters of the *lpmA* orthologs TattoneD and LoenenD (data not shown). Removal of this GC-rich motif from the 300 bp promoter of *lpmA* by site-directed mutagenesis resulted in a strong decrease in the ability of the promoter to drive dTomato expression (Figure 6C), since only weak fluorescence is observed. This confirms that the motif is indeed the binding site of Roc1 and that this binding site is required for correct promoter activity.

## DISCUSSION

Fungal deconstruction of lignocellulose in the plant cell requires the complex orchestration of a broad set of enzymes, and the expression of these enzymes is generally tailored to the type of polymers in the substrate by transcription factors. Several such transcriptional regulators have been identified in the phylum Ascomycota, but to date not in the phylum Basidiomycota. These phyla diverged over 600 million years ago^1^. Here, we identified the transcription factor Roc1 as a regulator of cellulase expression in the wood-decaying mushroom *Schizophyllum commune*. A *roc1* deletion strain cannot efficiently utilize cellulose (and, to a lesser extent, hemicellulose) as a carbon source. Moreover, ChIP-Seq revealed that Roc1 binds the promoters of various types of cellulase genes (including several lytic polysaccharide monooxygenases) while growing on cellulose, indicating that Roc1 directly regulates those genes. Furthermore, Roc1 activates its own expression, likely in a positive feedback loop.

*S. commune* is a highly polymorphic basidiomycete, both phenotypically and genetically. Strains H4-8, TattoneD and LoenenD varied considerably in their growth profiles, and showed a high variation in their genomes. This extraordinary genetic diversity was previously shown for other strains of *S. commune* as well^72^. Despite the high level of sequence variation between the strains, the number of CAZyme genes was remarkably similar. Therefore, it is challenging to link the phenotypical differences in growth profiles to a signature in the genome. However, the comparative transcriptomics approach allowed us to detect conserved responses in the expression profile, despite the high strain diversity. Although there is considerable variation between how the strains change their expression profile to the various carbon sources, the expression profile of CAZymes is more conserved. Furthermore, although the response of transcription factors was generally less conserved than the response of CAZymes, we identified a single transcription factor gene (*roc1*) that was consistently up-regulated under these conditions in both strains.

Cellulose, cellobiose, and xylan are major constituents of lignocellulose in wood. Cellulose is a polysaccharide of β-1,4-linked glucose residues. Cellobiose is a dimer of β-1,4-linked glucose residues and is an intermediate breakdown product of cellulose during enzymatic digestion. Xylan is a group of hemicelluloses consisting of a backbone of β-1,4-linked xylose residues. The observation that growth on these carbon sources is specifically affected, but not on other carbon sources, is a strong indication that Roc1 regulates the process of lignocellulose degradation. Growth on pectin, another constituent of lignocellulose in wood, is not negatively affected in Δ*roc1*. This indicates that Roc1 is likely not directly involved in pectinase expression.

It was previously shown that transcription factors involved in the regulation of polysaccharide degradation are generally poorly conserved between ascomycetes and basidiomycetes ^10^, and this is also the case for Roc1. Even within the basidiomycetes, Roc1 is only conserved in the class Agaricomycetes. The majority of fungi in this class are wood-degrading^5^, although it also includes mycorrhizal and plant pathogenic fungi (some of which have a partially saprotrophic lifestyle). Inversely, wood-degrading fungi are predominantly found in the class Agaricomycetes. This correlation between lifestyle (lignocellulose-degrading) and the presence of a Roc1 ortholog suggests that the function of Roc1 may be conserved in other members of the Agaricomycetes as well.

Regulators of cellulases were previously identified in the distantly related ascomycete *N. crassa* ^15^. It is important to note that these transcription factors (CLR-1 and CLR-2) are not orthologous to Roc1 of *S. commune*, nor are these two genes conserved in *S. commune*. Remarkably, however, the binding motifs of CLR-1 and Roc1 show a large degree of similarity. The binding motif of Roc1 (CCG-N-CGG) is part of the CLR-1 motif (CGG-N_5_-CCG-N-CGG). It is tempting to speculate that this is an example of convergent evolution, or that the binding motif predates either Roc1 or CLR-1. In the latter case, one transcription factor could have taken over the role of the other in an ancestor of ascomycetes and basidiomycetes. It should be noted, however, that binding motifs of regulators of the Zn_2_Cys_6_ transcription factor family frequently contain a CCG triplet, as is also the case for Gal4 of *S. cerevisiae*^67^.

Orthologs of Roc1 in *Agaricus bisporus* (a litter-degrading, mushroom-forming fungus) and *Dichomitus squalens* (a white rot, mushroom-forming fungus) display an expression profile that is very similar to the profile in *S. commune*. In *A. bisporus* the ortholog of Roc1 is protein ID 224213 (version Agabi_varbis_H97_2 of the genome annotation). Previously, whole-genome microarray expression data was published during growth on (glucose-rich) defined medium as well as on compost^73^. The expression of *roc1* was more than 10-fold higher when grown on compost (which contains large amounts of complex lignocellulosic carbon sources) than when grown on glucose-rich medium. Similarly, the ortholog of Roc1 in *D. squalens* is protein ID 920001 (version Dicsqu464_1 of the genome annotation). Previously, whole-genome expression data was published during growth on cellulose (avicel) and on cellulose in combination with glucose^74^. Expression was considerably higher (more than 6-fold) when grown on cellulose alone, when compared to when grown on a combination of glucose and cellulose. Although difficult to directly compare to our study, both these studies show that *roc1* in these fungi is down-regulated in glucose-rich medium, similar to the situation in *S. commune*.

Roc1 not only binds to promoters of cellulases, but also to promoters of several transcription factors. This suggests that Roc1 not only regulates lignocellulose degradation, but that it is also an important regulator of other downstream processes. While Roc1 ChIP-Seq revealed an enrichment of binding sites near lignocellulolytic CAZymes, the majority of putative binding sites were not in the promoter region of CAZymes or even genes up-regulated during growth on cellulose. It is currently not known what the role of Roc1 is in the regulation of these binding sites. A similar number of peaks has previously been reported for the Zn_2_Cys_6_ transcription factor PRO1 in *Sordaria macrospora*^75^, while fewer peaks were reported for other Zn_2_Cys_6_ transcription factors, including AflR in *Aspergillus flavus*^76^, CrzA in *Aspergillus fumigatus*^77^ and FgHtf1 in *Fusarium graminearum*^78^. Since the consensus sequence of the binding motif is rather short, it seems likely that not every occurrence of the motif results in a change of gene expression after binding by Roc1. Identifying binding sites during growth on additional substrates, in combination with RNA-Seq, may reveal more insights into the role of Roc1 in regulating lignocellulose degradation.

This is the first report of ChIP-Seq with a specific transcription factor in a mushroom-forming fungus. It is a crucial step to mapping the regulatory networks of transcription factors in this group of fungi, since direct interactions between transcription factors and promoters can now be revealed in vivo.

This will also be useful for studying the direct targets of the transcription factors involved in mushroom development^19,21,71,79^ and other processes in this important group of fungi.

## DATA AVAILABILITY

All genome assemblies and annotations can be interactively accessed through the JGI fungal genome portal MycoCosm^29^ at http://mycocosm.jgi.doe.gov. The data are also deposited in DDBJ/EMBL/GenBank under the following accessions numbers [*submission in progress, will be released upon publication*] for *S. commune* H4-8 (version Schco3), [*submission in progress, will be released upon publication*] for *S. commune* TattoneD (version Schco_TatD_1), and [*submission in progress, will be released upon publication*] for *S. commune* LoenenD (version Schco_LoeD_1).

The RNA Sequencing reads have been deposited in the NCBI Short Read Archive under project IDs SRP048482 (strain H4-8 on various carbon sources) and SRP053470 (strain TattoneD on various carbon sources). The ChIP-Seq reads have been deposited in the NCBI Short Read Archive under bioproject ID PRJNA726034.

## ACKNOWLEDGEMENTS

The work conducted by the U.S. Department of Energy Joint Genome Institute, a DOE Office of Science User Facility, is supported by the Office of Science of the U.S. Department of Energy under Contract No. DE-AC02-05CH11231. This project has received funding from the European Research Council (ERC) under the European Union’s Horizon 2020 research and innovation programme (grant agreement number 716132). We thank Utrecht Sequencing Facility for providing sequencing service and data for the ChIP-Seq analysis. Utrecht Sequencing Facility is subsidized by the University Medical Center Utrecht, Hubrecht Institute, Utrecht University and The Netherlands X-omics Initiative (NWO project 184.034.019). We thank Steven Ahrendt for technical assistance with data submission to GenBank.

## AUTHOR CONTRIBUTIONS

Performed experiments and analyzed the data: IMM, PJV, IDV, BB, AC, CD, HL, AL, HP, MBPS, MT, AT, JS, JG, LGG, RAO. Supervision/coordination of experiments: KB, JG, LGG, IGC, HABW, IVG, RAO. Wrote the manuscript: IMM, PJV, IDV, RAO. Provided funding: IGC, HABW, IVG, RAO. Conceived the project: RAO. Read and approved the manuscript: all authors

## SUPPLEMENTARY TABLES

**Table S1.** The species used in this study and their Roc1 orthologs. The previously published genome and genes were obtained from the indicated publications. The indicated phylogeny is based on the NCBI taxonomy database^80^.

**Table S2.** Primer used in this study.

**Table S3.** Statistics of the assemblies and gene predictions. For strain H4-8 an updated assembly and annotation were generated (version Schco3). Strains TattoneD and LoenenD were newly sequenced for this study.

**Table S4.** Genes encoding carbohydrate-active enzymes (CAZymes) in the three strains of *S. commune*.

**Table S5.** Expression values of genes (in RPKM) of strains H4-8 (first tab in the Excel file) and TattoneD (second tab) on medium containing either glucose, cellulose or wood as sole carbon source. Functional annotations of the genes are given. For each comparison of two growth conditions, three columns are given to describe the differential expression: the q-value (calculated by Cuffdiff), the log2 of the ratio of expression values (after increasing those with 1 to avoid division by zero issues), and whether or not the differential expression may be considered biologically relevant (indicated with ‘yes’ if the q-value is lower than 0.05, and the fold-change is at least 4, and at least one of the two conditions in question has an expression value of at least 10 RPKM).

**Table S6.** Locations of the peaks identified in the ChIP-Seq analysis. These peaks can be regarded as Roc1 binding sites. The genes associated with these peaks, as well as information about their annotation and expression profile are indicated.

**Table S7.** Enrichment of functional annotation terms among genes associated with a Roc1 binding site.

## SUPPLEMENTARY FIGURES

**Figure S1.**
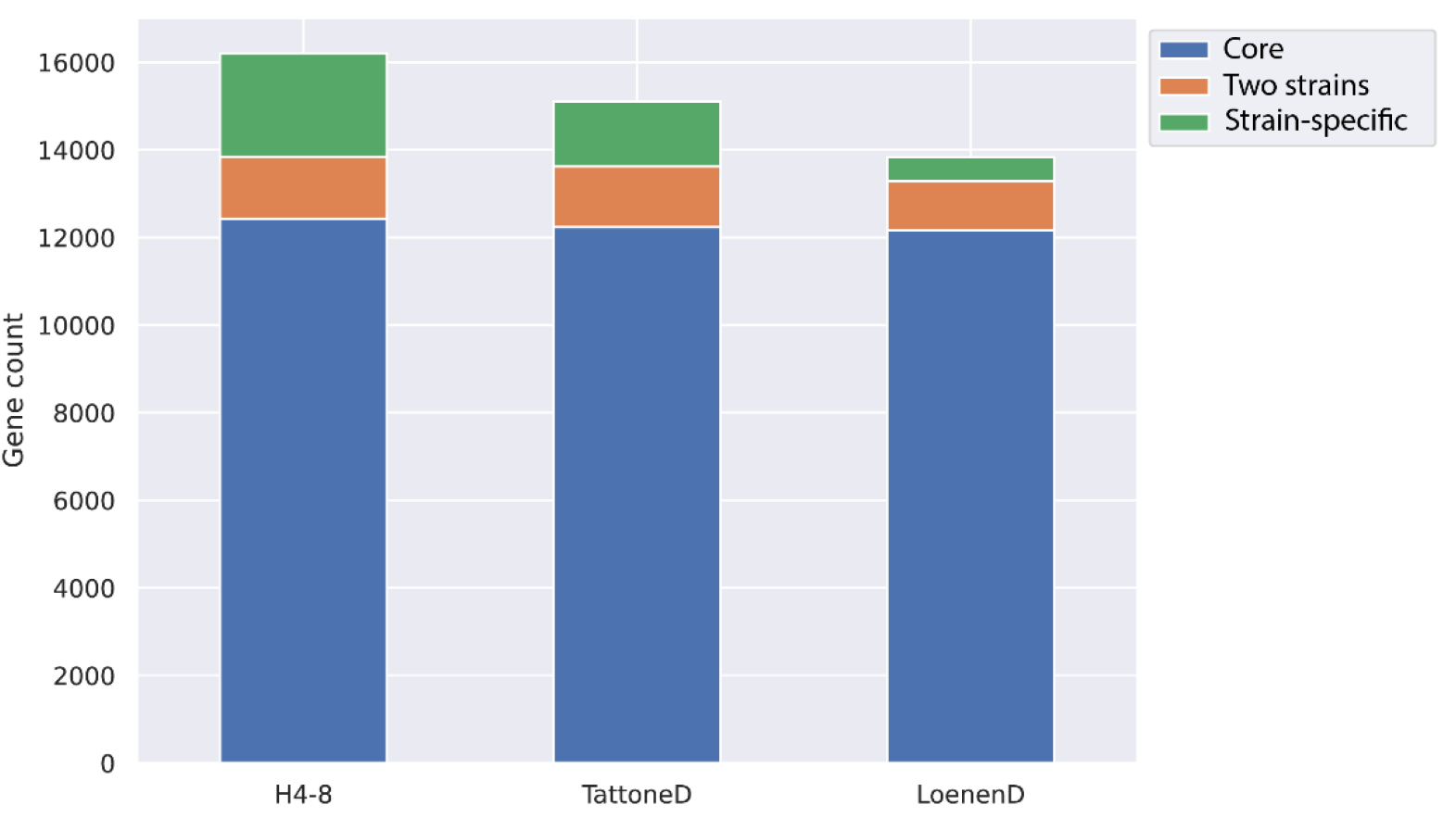
The number of predicted genes per strain and their conservation. The conservation was determined with Orthofinder. Genes in an orthogroup that had members from all three strains were labeled as ‘core’, genes in orthogroups that had members from two strains were labeled as ‘two strains’, and the remaining genes were labeled as ‘strain-specific’.

**Figure S2.**
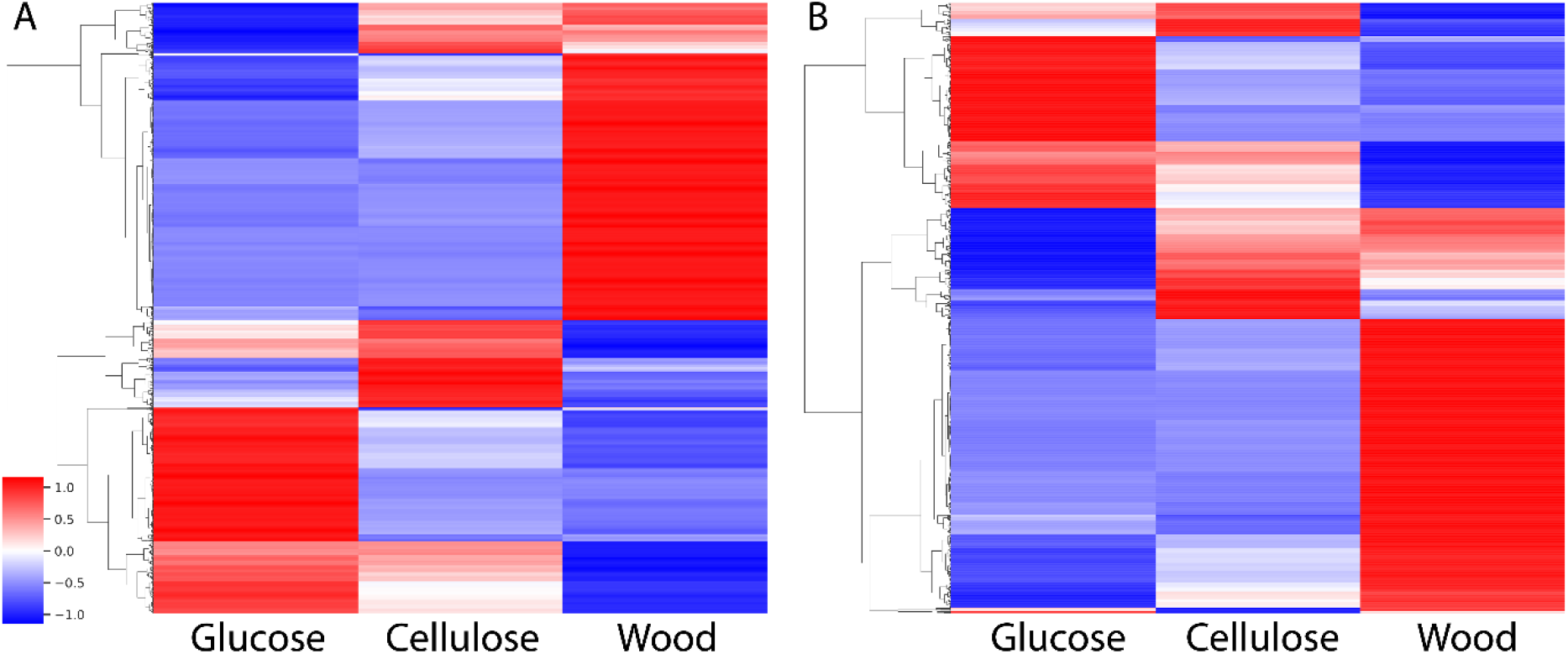
Heat map of gene expression during growth on either glucose, cellulose or wood as sole carbon source. Only genes that are differentially expressed between at least two conditions are depicted. The expression values were z-transformed, resulting in a z-score. All expression values are in Table S4. **A**. Strain H4-8. **B**. Strain TattoneD.

**Figure S3.**
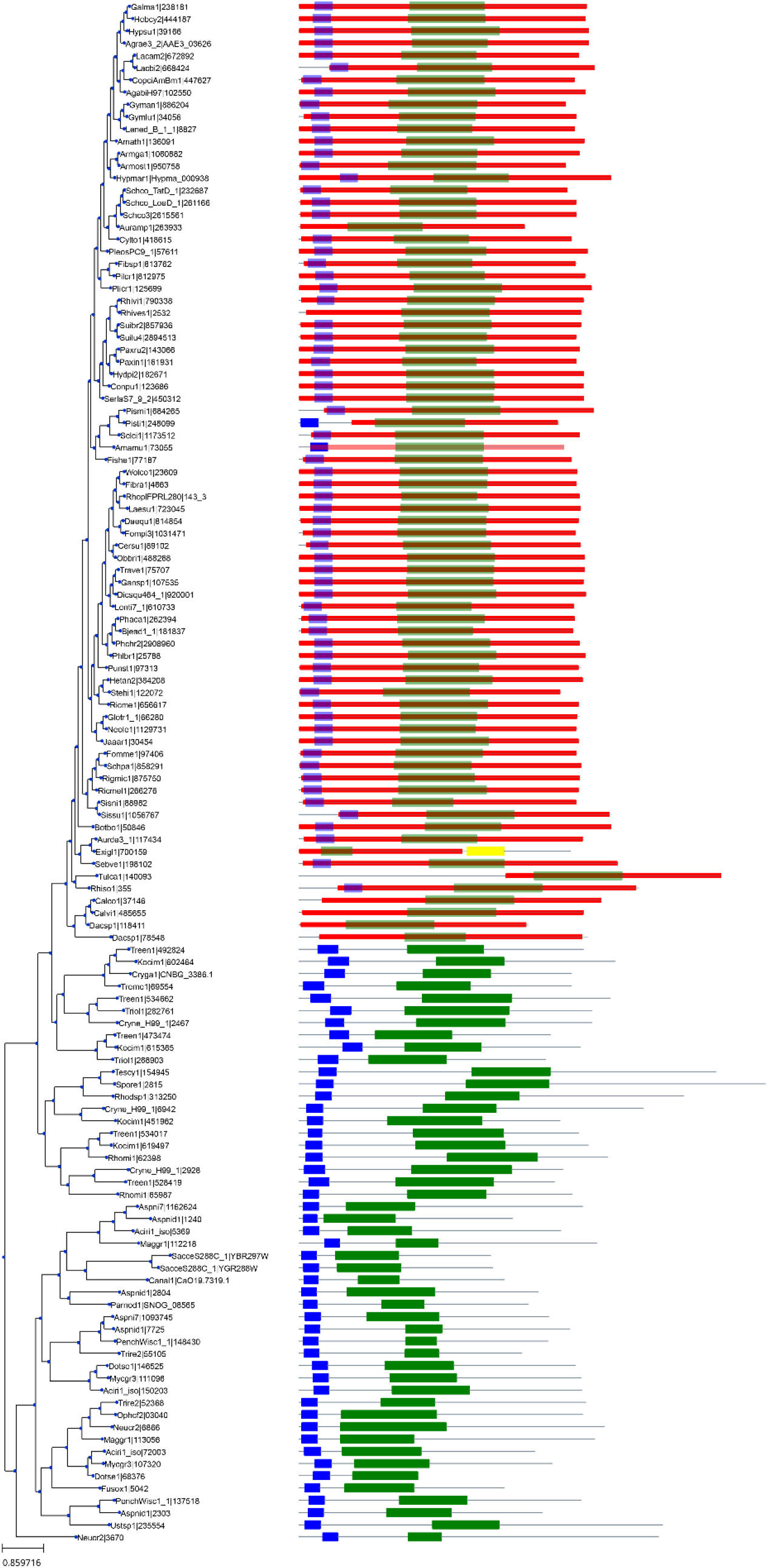
Gene tree of Roc1 orthologs and more distant homologs. The locations of the PFAM domains PF00172 (Zn_2_Cys_6_ binuclear cluster domain) and PF04082 (Fungal specific transcription factor domain) are indicated in blue and green, respectively. The location of a conserved Roc1 HMM domain is indicated in red. A protein is indicated as a Roc1 ortholog if the blue, green and red domains are present, whereas other proteins in this tree are considered more distant homologs. Branch lengths between the Roc1 orthologs are generally considerably shorter than the branch lengths between the more distant homologs. Roc1 orthologs are only found in Agaricomycetes (Figure S4). The first part of the protein IDs represents a species code (see Table S1 for the full species name).

**Figure S4.**
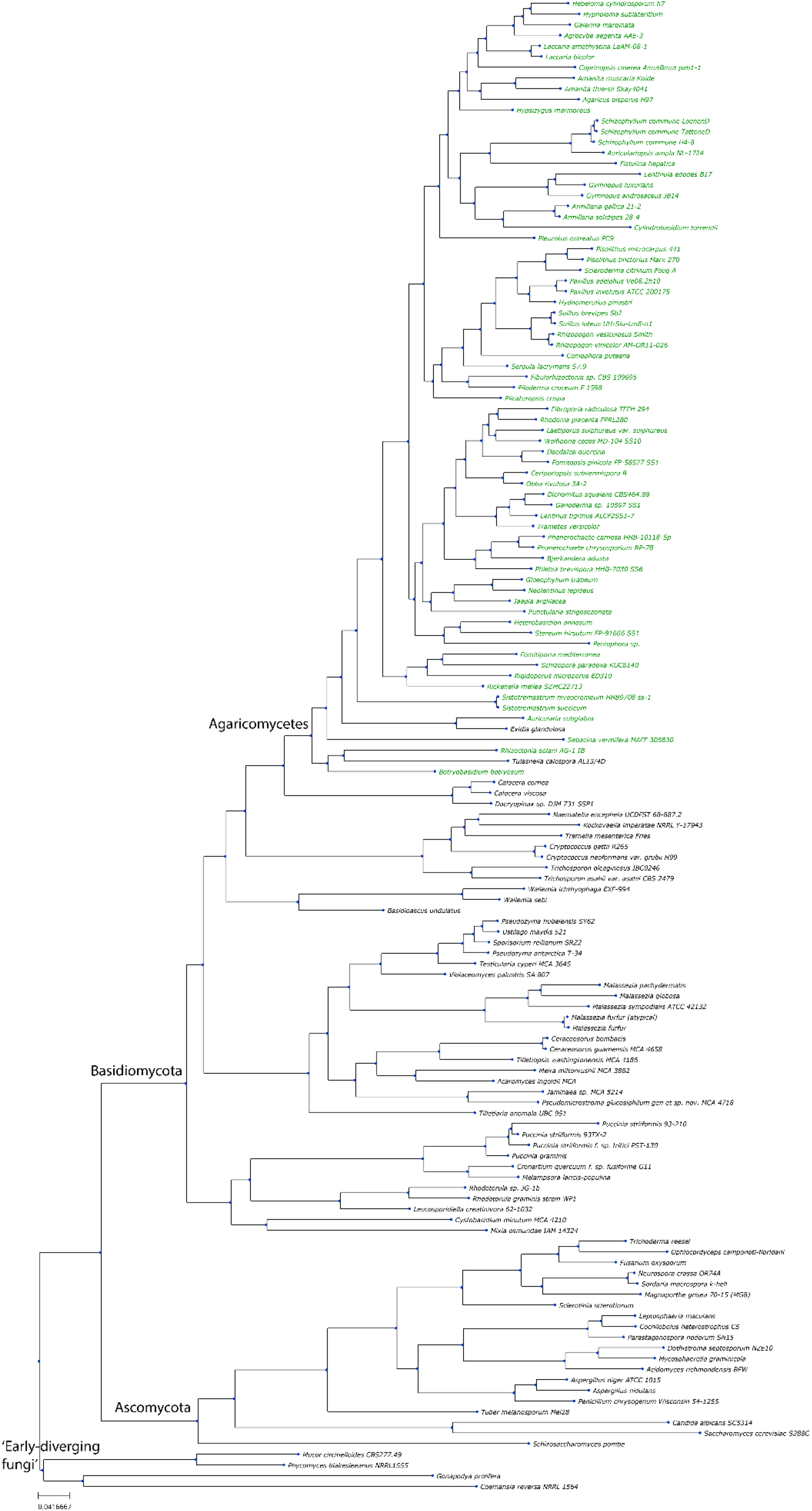
Species tree of 140 fungal species. The species with a putative Roc1 ortholog are indicated in green. Roc1 is only conserved in the Agaricomycetes. See Table S1 and Figure S3 for the protein IDs of the Roc1 orthologs.

**Figure S5:**
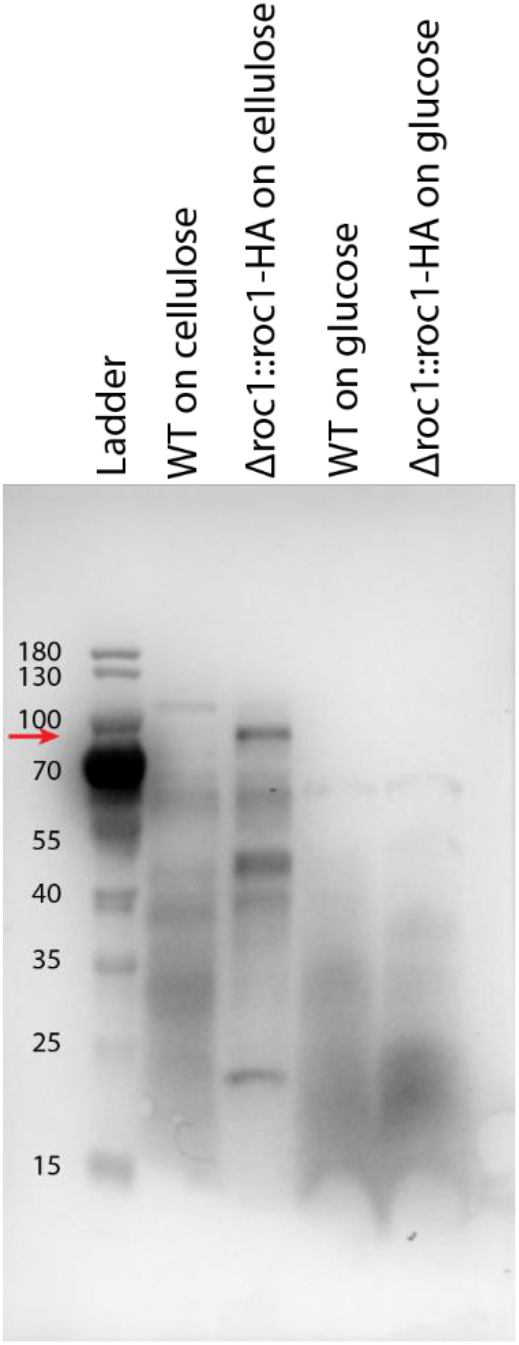
Western Blot confirms the expression of the HA-tagged Roc1 protein. The wild type (i.e. reference strain H4-8) and *Δroc1::roc1-HA* (i.e. the deletion strain complemented with the gene encoding the HA-tagged Roc1) were grown on cellulose or glucose. The predicted size of the HA-tagged Roc1 protein is 78.9 kDa. A band of this size is visible in the complemented deletion strain when grown on cellulose, but not when grown on glucose (the height of the band is indicated with a red arrow). As expected, this band is not found in the wild type strain under either condition.

**Figure S6.**
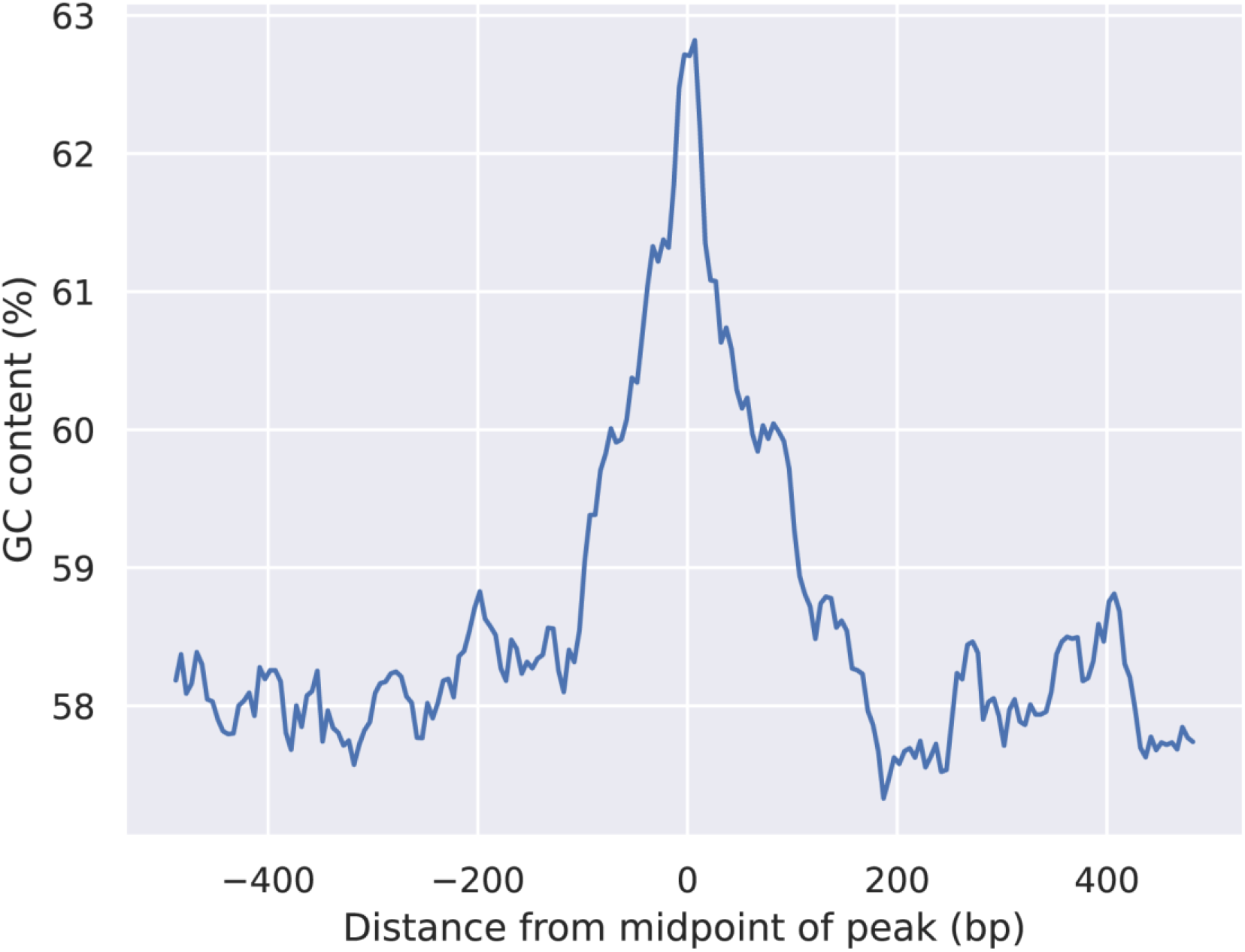
GC content along the length of the 1427 ChIP-Seq peaks. Around the midpoint of the peaks there is an increase of the GC content, which indicates that the Roc1 binding motif is GC-rich.

**Figure S7.**
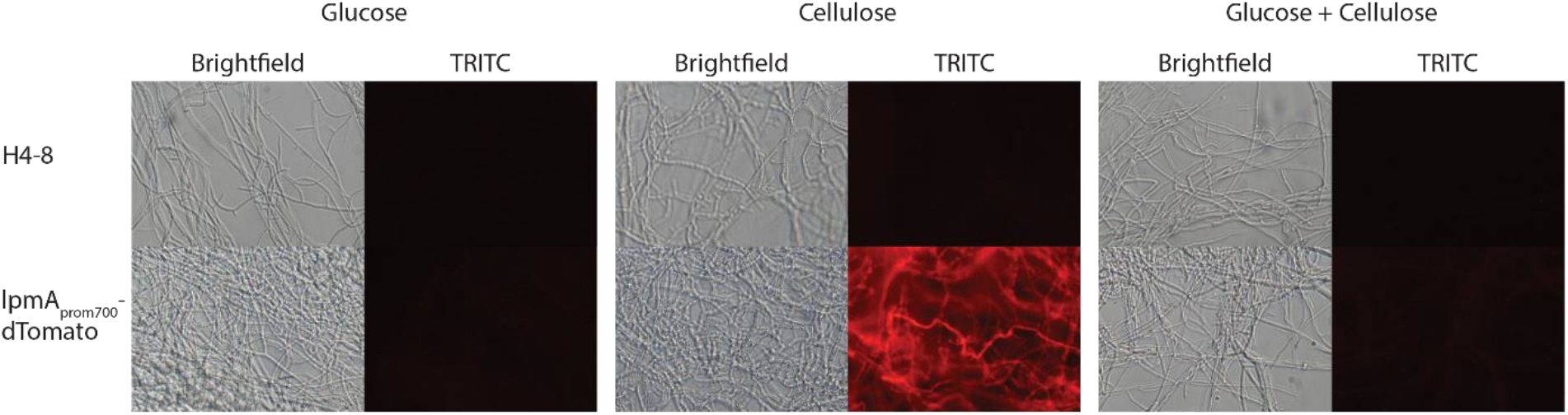
Fluorescence of dTomato driven by the 700 bp promoter of lpmA. When grown on glucose, no fluorescence is observed. In contract, when grown on cellulose strong fluorescence is observed, in concordance with the expression profile of lpmA. When grown on a mix of glucose and cellulose, no fluorescence is observed, indicating the carbon catabolite repression overrules the induction by cellulose. In the reference strain H4-8, no fluorescence is observed under any condition.

